# Transcriptomic splicing analysis reveals a gene-independent form of TE reactivation in tumors representing unrecognized source of oncogenes and neoantigens

**DOI:** 10.1101/2025.03.01.640928

**Authors:** Ziteng Li, Yichao Bao, Qin Li, Hongwu Yu, Hena Zhang, Yifan Wen, Yu Yang, Jingjing Zhao, Peng Lin, Yan Li, Zhixiang Hu, Xichun Hu, Xiaodong Zhu, Shenglin Huang

**Affiliations:** Department of Integrative Oncology, Fudan University Shanghai Cancer Center, and Shanghai Key Laboratory of Medical Epigenetics, International Co-laboratory of Medical Epigenetics and Metabolism, Institutes of Biomedical Sciences, Shanghai Medical College, Fudan University, Shanghai, China; Department of Medical Oncology, Fudan University Shanghai Cancer Center, Shanghai, China; Department of Oncology, Shanghai Medical College, Fudan University, Shanghai, China

**Author notes:** These authors contributed equally to this work (Ziteng Li, Yichao Bao). **Correspondence**: Shenglin Huang; Xiaodong Zhu; Xichun Hu.

## Abstract

Transposable elements (TEs) are reactivated in tumors and serve as significant contributors to tumor transcriptomic complexity and heterogeneity. However, their repetitive nature and diverse transcriptional forms have hindered efforts to characterize TE-derived transcripts independent of host gene contexts. Here, using our self-developed splicing-junction analysis tool ASJA, we systematically interrogated TE transcription across 32 cancer types, identifying transcripts autonomously initiated from and exclusively spliced among TEs with high TE content, termed TEtrans. Our analysis revealed 5,361 TEtrans exhibiting pan-cancer prevalence, tumor-specific expression and features of mature transcripts, including canonical splicing, 5’ caps and polyA tails. Mostly unannotated and enriched in intergenic regions, TEtrans are predominantly derived from primate-specific classes with conserved splice pattern. TEtrans burden demonstrates heterogeneous associations with prognosis and immune activity across cancers, while their expression can be epigenetically dual-regulated by stemness- and inflammation-associated transcription factors. Functional studies uncovered that TEtrans could act as oncogenes, exemplified by tsTE1, a HERVH-derived transcript that promotes colorectal cancer proliferation by enhancing TOP1-mediated DNA supercoil relaxation. Remarkably, ∼7.2% of TEtrans are identified to encode tumor neoantigens, including viral proteins and unannotated peptides, which are shared among patients and validated by proteogenomic analysis. These TEtrans-derived neoantigens are immunogenic both in vitro and in vivo, and exceed neoepitopes from genomic alterations in abundance per tumor, particularly in cancers with low mutational burden. Collectively, as a gene-independent form of TE-derived transcripts, TEtrans represents a unique source of oncogenes and tumor neoantigens, expanding the functional repertoire of TEs in cancer biology and offering new avenues for therapeutic targeting.

**Highlights:**

1. TEtrans are tumor-specific transcripts with conserved splicing patterns, predominantly derived from primate TEs in intergenic regions.
2. TEtrans expression is epigenetically activated and dually regulated by stemness/inflammation transcription factors.
3. TEtrans function as oncogenes, exemplified by tsTE1 enhancing TOP1-mediated DNA relaxation.
4. TEtrans encode tumor neoantigens that elicit CD8+ T cell responses, providing targets for immunotherapy, especially in low-mutation tumors.

**Abstract figure:** 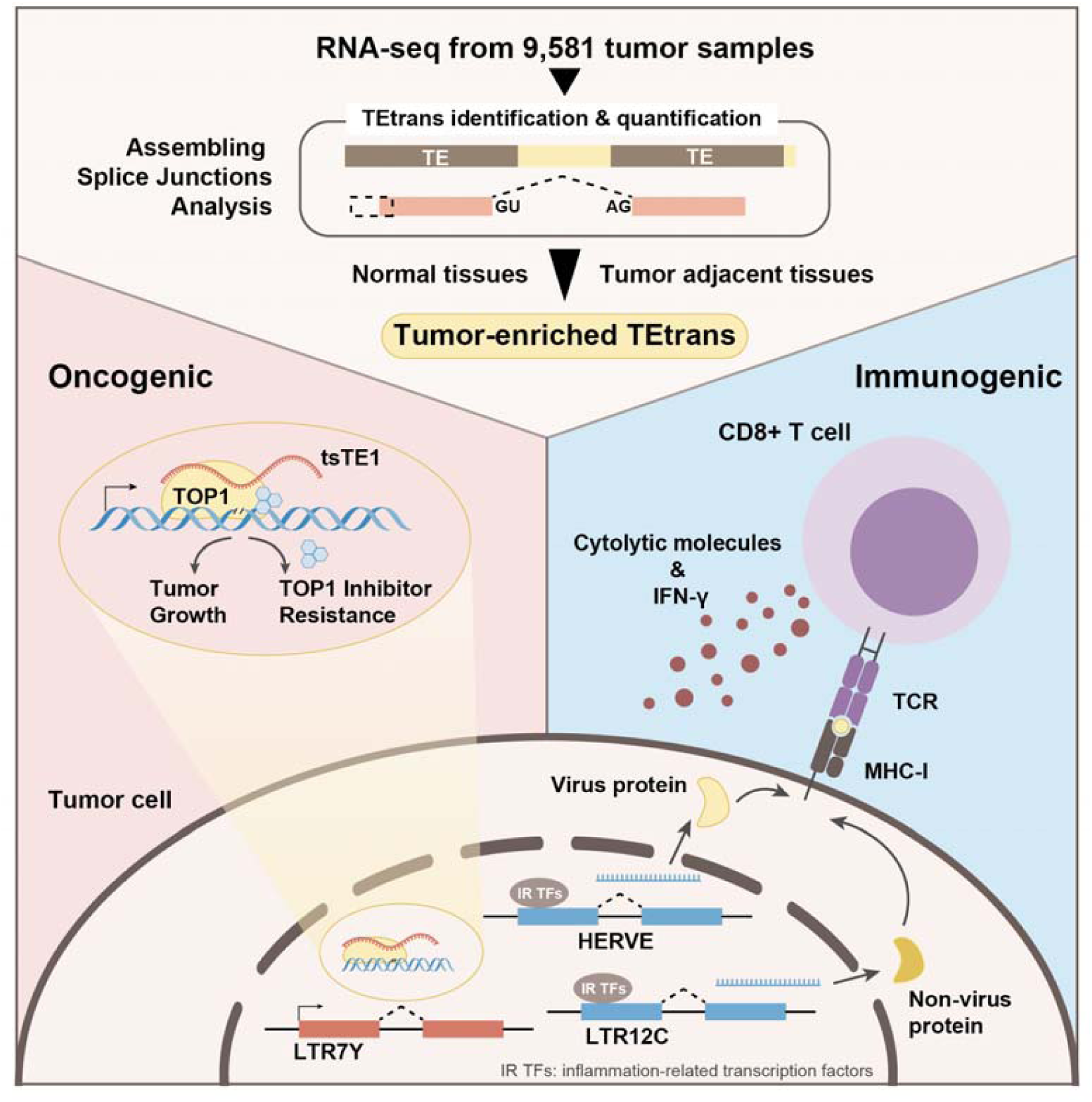

## Introduction

Tumorigenesis is widely recognized as a complex process driven by the accumulation of genetic and epigenetic alterations that disrupt normal gene expression profiles ^1^. Although nearly three-quarters of the human genome can be transcribed, only a small fraction (approximately 2%) of RNA sequences are derived from protein-coding genes ^2^. This discrepancy underscores the importance of exploring the non-coding transcriptome, particularly in the context of cancer, where aberrant transcriptional activity can contribute to tumor progression and heterogeneity. A deeper understanding of the tumor transcriptome’s complexity is crucial for identifying novel diagnostic and therapeutic targets, as well as for developing innovative treatment modalities.

Transposable elements (TEs), which constitute approximately half of the human genome, are mobile DNA sequences that have historically been dismissed as "junk DNA" ^3,4^. These elements were primarily integrated into the host genome through viral infections and other evolutionary processes, with most having lost their ability to transpose over time ^5^. Under normal physiological conditions, TE transcription is tightly regulated and suppressed by mechanisms such as DNA methylation ^6,7^, transcription factors ^8–10^, and PIWI-interacting RNAs ^11^. However, in cancer, epigenetic dysregulation, particularly widespread hypomethylation, leads to the reactivation of numerous silent TEs. This reactivation contributes to the unique complexity and diversity of tumor transcriptome ^6,7,12^.

TE transcripts play a dual role in cancer: they can promote tumor malignancy while also serving as potential targets for immune recognition ^12^. TE-gene chimeric transcripts and TE-derived viral proteins can act as oncogenes, boosting tumor proliferation, while simultaneously generating novel epitopes that are presented by Major Histocompatibility Complex (MHC) molecules, thereby activating T cell-mediated immune responses ^13–16^. Epigenetic drugs, such as DNA methyltransferase inhibitors (DNMTis) and Histone Deacetylase inhibitors (HDACis), have been shown to potentiate TE transcription, leading to the accumulation of cytoplasmic double-stranded RNAs (dsRNAs) which can trigger the viral mimicry responses ^5,17–19^. Thus, TE transcriptional activation not only provides oncogenic targets for therapeutic intervention but also offers a promising avenue for enhancing anti-tumor immunity. Despite the profound translational potential of TE-derived transcripts, their repetitive nature and complex transcriptional forms pose significant challenges for accurate localization and quantification, hindering a comprehensive understanding of their roles in cancer ^20^.

Current research on TE transcription often quantifies TEs at the family level, which obscures the contributions of individual TE copies with distinct genomic locations ^6,21,22^. Alternatively, studies may emphasize on TE-gene chimeric transcripts while underestimating the significance of TE-derived RNAs originating from intergenic regions ^15,16,23^. In fact, TE-derived RNAs can include gene-independent products that undergo post-transcriptional processing, such as 5’ capping, splicing, and 3’ polyadenylation. Some of these transcripts encode complete or partial proviruses, such as syncytin-1 and syncytin-2, which play roles in syncytialization during embryogenesis. Numerous intergenic HERV loci have been found to be expressed in normal tissues, though many result from read-through transcription. Notably, our previous work has identified tumor-specific activation of intergenic TEs based on RNA splicing, leading to the generation of gene-independent transcripts that promote cancer progression ^24,25^. These findings highlight the potential of TEs to autonomously generate mature, gene-independent transcripts that may contribute to tumorigenesis, warranting further investigation at the locus-specific level. Such transcripts, likely widespread in tumors, have not yet been systematically characterized to our knowledge.

In this study, we employed our previously developed RNA splicing identification and quantification tool, ASJA (Assembling Splice Junctions Analysis), which is designed to identify, extract, and quantify RNA splice sites from RNA-seq data ^26^. ASJA has proven highly effective in uncovering tumor-specific transcriptomes, enabling the discovery of novel transcripts that evade conventional annotation pipelines^24,25,27,28^. By localizing and quantifying splice sites, we can distinguish transcripts derived from different TE copies, enabling a site-specific comparison of these transcripts and the selection of tumor-specific TE-derived ones. We systematically identified and analyzed TE transcription, focusing on transcripts initiated from and exclusively spliced among TEs with high TE content. We defined this unique class of TE-derived transcripts as TEtrans. Leveraging the advantages of ASJA, we comprehensively characterized TEtrans in tumors, elucidating their origins, splicing patterns and regulatory mechanisms, while also revealing their dual functional roles as oncogenes and immune vulnerabilities. Our findings not only delineate a previously neglected dimension of TE transcription in cancer but also establish TEtrans as a novel source of tumor-specific targets and antigens, holding promise for the development of innovative cancer therapies, including personalized vaccines and immunotherapies.

## Results

### TE splicing events help identify novel form of TE transcription with recurrent and tumor-enriched expression

We used ASJA program to extract and quantify assembled transcript junctions from RNA sequencing data across TCGA tumor and adjacent samples, normal tissues from GTEx and tumor cell lines from CCLE **(Fig. 1A)**. Then we comprehensively assessed splicing events with donor or acceptor splice sites contributed by TEs, identifying 149,269 TE-associated splicing events from TCGA pan-cancer samples while only 40,737 from GTEx normal tissues excluding testis **(Fig. S1A)**. Over 86% TE splicing events detected in tumor tissues were simultaneously shared by tumor cell lines **(Fig. S1A)**. Annotating the genomic locations of splicing junctions, we showed that both donor and acceptor splice sites were more enriched in intergenic and intronic regions in contrast to all splicing events **(Fig. S1B)**. These observations verified the reactivation of TEs in tumors meanwhile suggested that TE splicing events are mainly driven by tumor cells and derived from non-coding regions.

**Figure 1:**
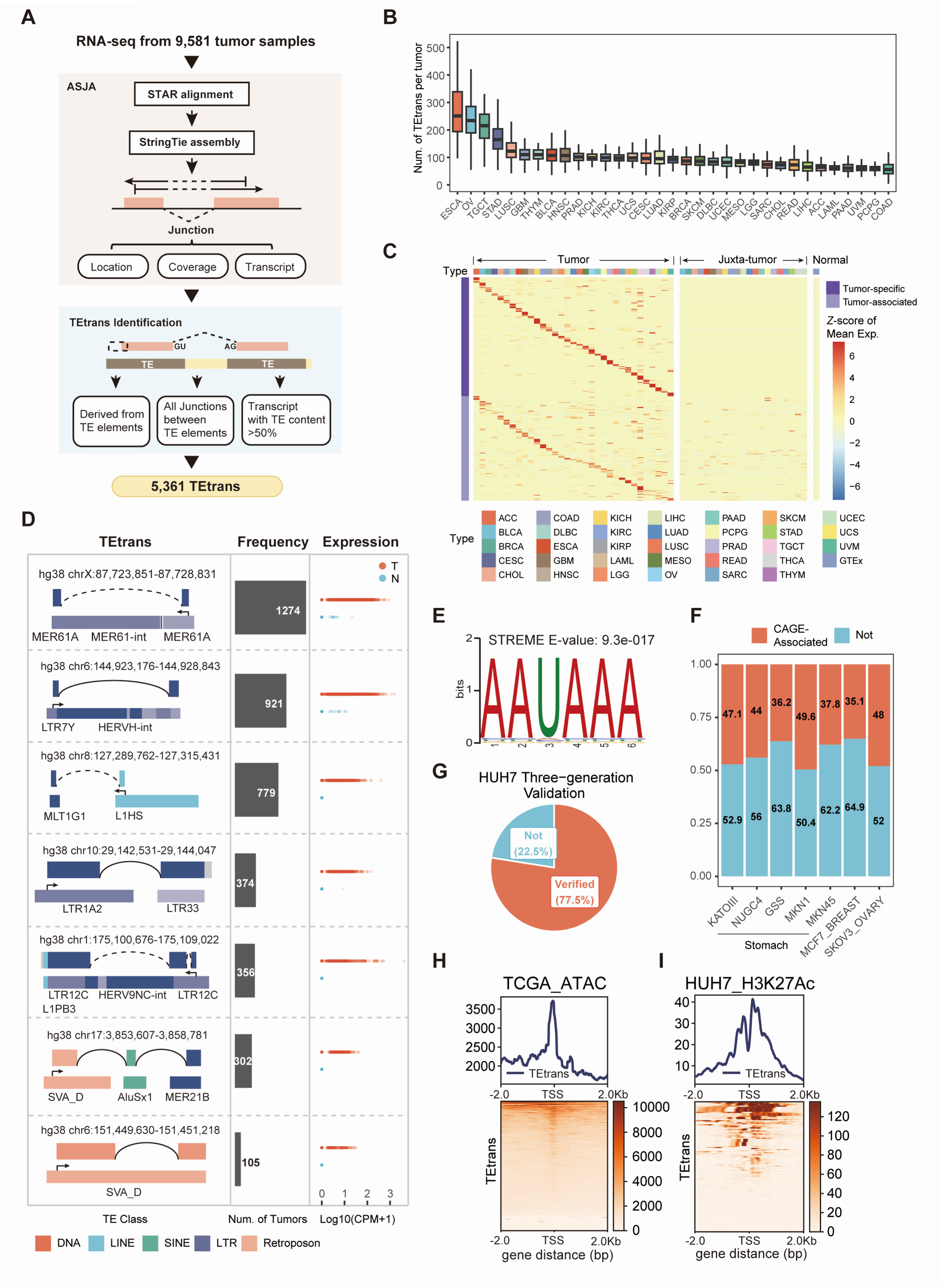
Identification and characterization of TEtrans, a gene-independent class of TE-derived and -spliced transcripts enriched in tumors. **A.** The TEtrans identification workflow. The ASJA program was employed to extract and quantify TE splicing events from RNA sequencing data of 9,581 TCGA tumor samples. Based on these splicing junctions, TEtrans was identifed as transcripts originating from TEs, with all splice sites occurring between TEs and containing more than 50% TE content, which consequently obtained 5,361 transcripts. **B.** Distribution of TEtrans across different tumor types in TCGA. **C.** TEtrans exhibits high tumor specificity. The heatmap shows the expression levels of the top 10 most expressed tumor-specific or associated TEtrans across various tumor types, demonstrating lineage-specific expression patterns. The heatmap colors represented the Z-score of mean expression for each TEtrans. **D.** Typical examples of TEtrans prevalent in tumor samples. The leftpanel shows the diagram of the transcript with the TE-originating and -splicing features of TEtrans illustrated. Genomic locations and TE subfamilies were annotated. Color schemes of box represent different TE class whereas positive or negative strand were denoted by solid or dashed lines respectively. The middle panel shows the number of tumor samples each TEtrans is expressed in. The rignt panel displays the expression of each TEtrans in tumor and adjacent samples. The TEtrans candidates showed are only detected in tumor samples or expressed five fold higher in tumor than adjacent tissues, demonstrating tumor-specific expression pattern. **E.** Significant enrichment of AAUAAA polyadenylation signals at the 3’ end of TEtrans transcripts, with an E-value representing the adjusted p-value. **F.** CAGE peak signals were detected within 500bp upstream and downstream of the TSS of TEtrans. CAGE-related TEtrans accounts for the range of 35.1%-49.6% in gastric, breast, and ovarian cancer cell lines. **G.** Validation of TEtrans splicing sites using third-generation sequencing data from HUH7 cell lines. **H-I.** Enrichment of transcriptional activation signals near the promoter regions of TEtrans in TCGA tissue samples and HUH7 cell lines. (H) shows ATAC-seq peak signal enrichment near TEtrans TSS in TCGA samples, while (I) shows the enrichment of H3K27Ac peak signals near TEtrans TSS in HUH7 cell lines.

Previous studies on TE splicing events are mostly related to gene regions, such as TE-derived alternative splicing isoforms whereas those located in intergenic regions as identified by ASJA were possibly neglected. A TE-derived transcript derived and spliced from intergenic TEs was unexpectedly identified while investigating tumor-specific transcripts in our previous work^25^, leading to the speculation that TEs may autonomously transcribed and spliced independent of genes, a phenomenon likely specific to tumors. Interrogating transcription start site (TSS), splice sites as well as transcript sequences, we obtained 5,361 transcripts initiated from and exclusively spliced among TEs with high TE content, namely TEtrans, from 9,581 TCGA pan-cancer tumor samples **(Fig. 1A)**. TEtrans contained substantial novel transcripts, with approximately 91.7% splicing events involved in TEtrans are unannotated **(Fig. S1C)**.

TEtrans could be detected in almost all TCGA tumor samples (8969/8976, 99.92%) and all of CCLE tumor cell lines, with a median number of 89 TEtrans per tumor tissue and 146 per cell line **(Fig. 1B, Fig. S1F)**. Upper gastrointestinal tumors including esophageal cancer (ESCA) and gastric cancer (STAD), reproductive system tumors such as ovarian cancer (OV) and testicular cancer (TGCT), as well as lung squamous cell cancer (LUSC) were found with highest TEtrans abundance **(Fig. 1B)**. TEtrans showed tumor-enriched manner in both frequencies and expression level **(Fig. 1C, Fig. S1E)**. Over 80% TEtrans are exclusively expressed in tumor samples (Tumor-specfic) or at least 5-fold higher than all juxta-tumor and normal tissue samples (Tumor-associated) **(Fig. 1C)**. Certain TEtrans, especially those in the reproductive tumors, were universally detected across all tumor types whereas others displayed lineage-specfic expression **(Fig. S1G)**. We examined the transcript structure and expression of top prevalent TEtrans examples **(Fig.1D)**, with two of them validated in tumor cell lines by real-time qPCR **(Fig.S2A)**. The recurrent and tumor-enriched nature of these transcripts further support that TEtrans as an uncharacterized source of potential tumor targets.

To verify the validity of TEtrans, we subsequently explored the characteristics of TEtrans. Motif enrichment over 3’ end sequence of TEtrans revealed the presence of conserved motif AAUAAA signaling for polyadenylation **(Fig. 1E)**. The proportion of TEtrans enriched with this motif was comparable to those of mRNAs and lncRNAs **(Fig. S2B)**. Annotated CAGE peaks acquired from FANTOM5 project could be found in TSS±500bp of 20.5% of TEtrans **(Fig. S2C-D)**. This proportion increased to near 50% while specifying samples to certain tumor cell lines **(Fig. 1F)**. Such findings reflected that TEtrans serve as a class of mature transcript with canonical 5’ capping and polyA tails. Chromatin state and histone modifications around TSS were further interrogated to verify the epigenetic activation of TEtrans. The ATAC-seq data from 655 TCGA tumor samples of 22 tumor types and H3K27Ac ChIP-seq data of HUH7 cancer cell line both demonstrated strong enrichment of peak signals near the TSS regions of TEtrans in the metaprofile **(Fig. 1H-I)**, indicating the accessible chromatin state for the transcription of TEtrans. Additionally, considering the short read length of RNA-seq and the high sequence repeatability of TEs, we further validated the splicing events of TEtrans in third-generation long-read RNA sequencing data. For HUH7 cell line, 77.5% splicing events of TEtrans were verified by full-length RNA sequencing **(Fig. 1G)**, proving the robustness of ASJA to identify TEtrans from the prospective of TE splicing. Collectively, these findings support the existence of TEtrans and further exhibited their mature nature harboring canonical transcription initiation signals and undergoing typical post-transcription processing.

### Examining splicing events uncovered the conservation of TEtrans splicing patterns

Since TEtrans were identified based on TE splicing events, we consequently investigated features of these splicing junctions. Locations of both donor and acceptor splice sites of TEtrans showed higher intergenic enrichment compared to the whole set of transcripts and all TE-derived splicing events **(Fig. 2A)**. Such intergenic proportion was increased to approximately 60% when only considering TEtrans specifically expressed in tumors. Analyzing TE classes, we observed that two TE classes, LTR and Retroposon, contributed more TE splicing sites than others, accounting for about 74% of the total TEtrans splicing events (TEtransjunc), which was also a marked increase in contrast with all TEs annotated by RepeatMasker (RM) as well as all spliced TEs identified (All TEjunc) **(Fig. 2B)**. TEs spliced in TEtrans harbor younger evolutionary ages than all TEs and most of them are primate-specific, consistent with the prominent enrichment of SVA and ERV families **(Fig. 2C-D)**. Besides, donor splice sites were additionally enriched in LINE1 family, which may be due to the promoter activities retained by some LINE1 elements.

**Figure 2:**
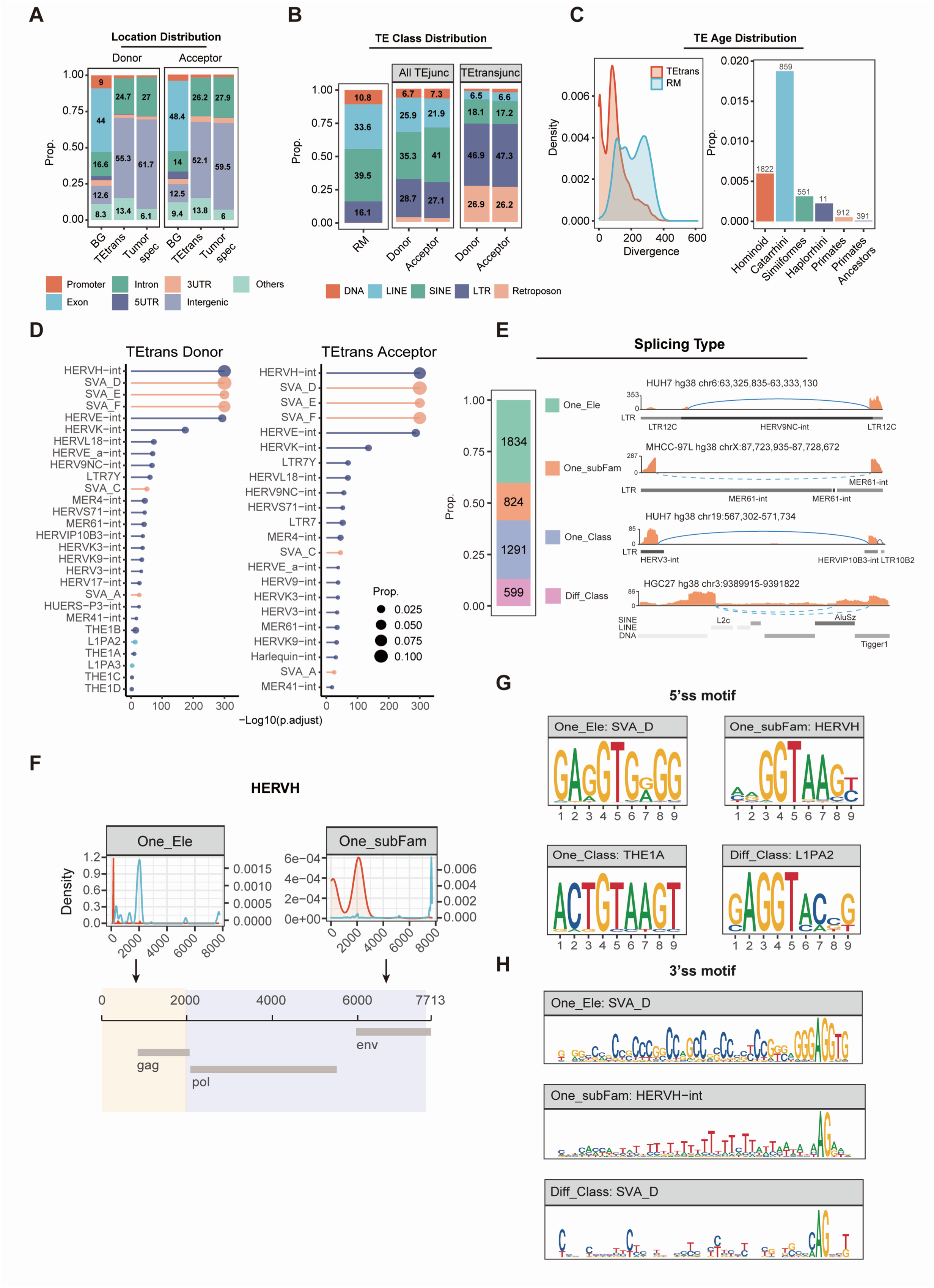
Examining splicing events uncovered the conservation of TEtrans splicing patterns enriched in primate-specific and evolutionarily young TEs. **A.** Genomic distribution of donor and acceptor splice sites within TEtrans. BG represents splice sites of all transcripts, and Tumor_spec refers to splice sites for tumor-specific TEtrans. **B.** Distribution of TE classes involved in TEtrans splicing. RM represents all TEs annotated by RepeatMasker, AllTEjunc refers to splice sites with at least one end derived from TE, and TEtransjunc indicates the splice sites specific to TEtrans. **C.** TEtrans splicing events predominantly occur in primate-specific and evolutionarily young TEs. **D.** Donor and acceptor splice sites in TEtrans are highly enriched in the ERV1 and SVA families, with donor sites also enriched in the LINE1 family. **E.** Proportions of the four splicing modes in TEtrans, along with representative schematic structures for each mode. TE classes and subfamilies are annotated in the examples, with different evolutionary ages represented by varying shades of gray. **F.** Upper panel: Donor and acceptor splicing positions for HERVH element in the One_Ele and One_subFam splicing modes, mapped to the consensus sequence of HERVH. Lower panel: Schematic representation of splicing position. The yellow part represents the One_Ele mode marking the gag region spliced out, whereas the purple region represents the One_subFam mode, highlighting the loss of pol and env ORFs. **G.** Examples for motifs of conserved 5’ splice sites (5’ss) in each splicing mode. 5’ss defined as a 9-mer motif extending 3bp upstream and 6bp downstream of the donor site. **H.** Examples for motifs of conserved 3’ splice sites (3’ss) for different splicing modes. 3’ss defined as a 43-mer motif extending 40bp upstream and 3bp downstream of the acceptor site. The SVA_D in One_Ele mode has a higher GC content at 3’ss compared to the Diff_Class mode.

As TEtrans were composed of a high fraction of TE sequences, TE sequences and splicing locations would significantly affect the ability of encoding retroviral protein, determining the potential function of TEtrans.Therefore, we first divided TEtrans splicing events according to hierarchical TE classifications and evolutionary ages into four patterns **(Fig. 2E)**: One_Ele refers to splicing within the same TE loci, some of which employs the canonical splice site of the element; One_SubFam denotes splicing between copies of the same element but with different ages, part of which could be generated by adjacent transposition during evolution; One_Class splicing occurred between distinct TEs of the same class, while Diff_Class refers to splicing between different TE classes. Of TEtrans splicing events, 40% displays One_Ele pattern and 87% occurs within the same TE classes, indicating that TE splice sites tend to be generated under similar sequence background **(Fig. 2E)**. Furthermore, we found that these four splicing patterns had distinct TE specificity: One_Ele, One_SubFam and One_Class were enriched in Retroposon, LTR and SINE class resepctively; LINE and DNA class preferred to spliced with other classes producing Diff_Class splicing **(Fig. S3A)**. Such results suggest that the splicing pattern preference of TEtrans may be attributed to the sequence difference.

The consensus sequence of a certain TE subfamily represented the average sequence of TE copies generated through multiple alignment and curation. We mapped the TE-contributed splice sites to the consensus sequence respectively and found that for certain TE subfamily, both donor and acceptor splice sites were highly conservative within the same splicing pattern whereas they may vary among different splicing types **(Fig. 2F, Fig. S3B)**. For example, HERVH elements belong to the endogenous retrovirus class 1 (ERV1) family which encodes viral proteins gag, pol, and env functioning as capsid, reverse transcriptase and envelope respectively for viral particles. Pol and env tend to be retained in One_Ele splicing yet more spliced in One_SubFam pattern, resulting in TEtrans with different viral protein coding frames thereby potentially exerting distinct functions **(Fig. 2F)**.

Enlightened by the repetitive nature and the concentrated junction breakpoints of TEs composing TEtrans, we further identified the conserved motifs for both donor (−3bp to +6bp for 5ss) and acceptor splice sites (−40bp to +3bp for 3ss). TEtrans splicing conforms to the classical GT-AG splicing rule. TEs demonstrated highly conserved 5ss in their preferred splicing modes, which was possibly attributed to less sequence divergence due to younger age of these TEs **(Fig. 2G, Fig. S3B).** We noted a clear discrepancy in the base compositions of 3ss motifs presumably containing polypyrimidine tract (PPT) for different TE derivations and splicing settings. Higher GC content was identified in 3ss of SVA_D element spliced in One_Ele way whereas PPT mainly composed with T bases was found in that of HERVH element spliced in One_subFam way **(Fig. 2H)**. To verify these splice sites, we paired the conserved 5ss and 3ss motifs derived from top enriched TEs, combined them into artificial introns and detected their splicing rate in 293T cell lines **(Fig. S3C)**. Confirming the authenticity of the identified TEtrans splice sites, most introns were spliced out at the ratio of over 50%, except the SVA family in One_Ele mode which was spliced less than that in Diff_Class pattern as well as other families **(Fig. S3D-E)**. As T content in the PPT was reported to be closely related to splicing efficiency ^29,30^, the low cleavage efficiency of the SVA One_Ele intron might be due to the high GC content of its 3ss. These splicing features of TEtrans may suggest the potential transcript functions and tumor-specific splicing mechanisms, representing another dimension of TE contributions beyond gene alternative splicing.

### Tumor TEtrans burden was heterogeneously associated with prognosis and immune microenvironment across tumor types

To measure the transcriptional load of TEtrans in tumor samples, we constructed TEscore based on the TEtrans expression levels and frequencies and then evaluated it in TCGA tumor samples **(Fig. 3A, Methods)**. We assessed the association of TEscore with patient prognosis and found that TEscore was significantly associated with at least one of four survival outcomes including OS, DSS, PFS, and DFS across 8 tumor types **(Fig. 3B)**. However, such prognostic associations can be opposite in different lineage context: high TEscore was a risk factor in hepatocellular and colorectal carcinoma, while was protective in cervical carcinoma and uveal melanoma **(Fig. 3B-C)**. Associations between TEscore and the immune microenvironment also varied among cancer types **(Fig. 3E)**. High TEtrans burden was linked to enhanced anti-tumor immunity in esophageal squamous cell carcinoma, breast cancer, and sarcoma, whereas it correlated to immune exhaustion in pancreatic, gastric, bladder, lung squamous cell, and head and neck squamous cell carcinomas. These finding implies that TEtrans may represent a mixed population of transcripts with divergent functions across different tumor species, highlighting the need for in-depth functional explorations over individual transcripts.

**Figure 3:**
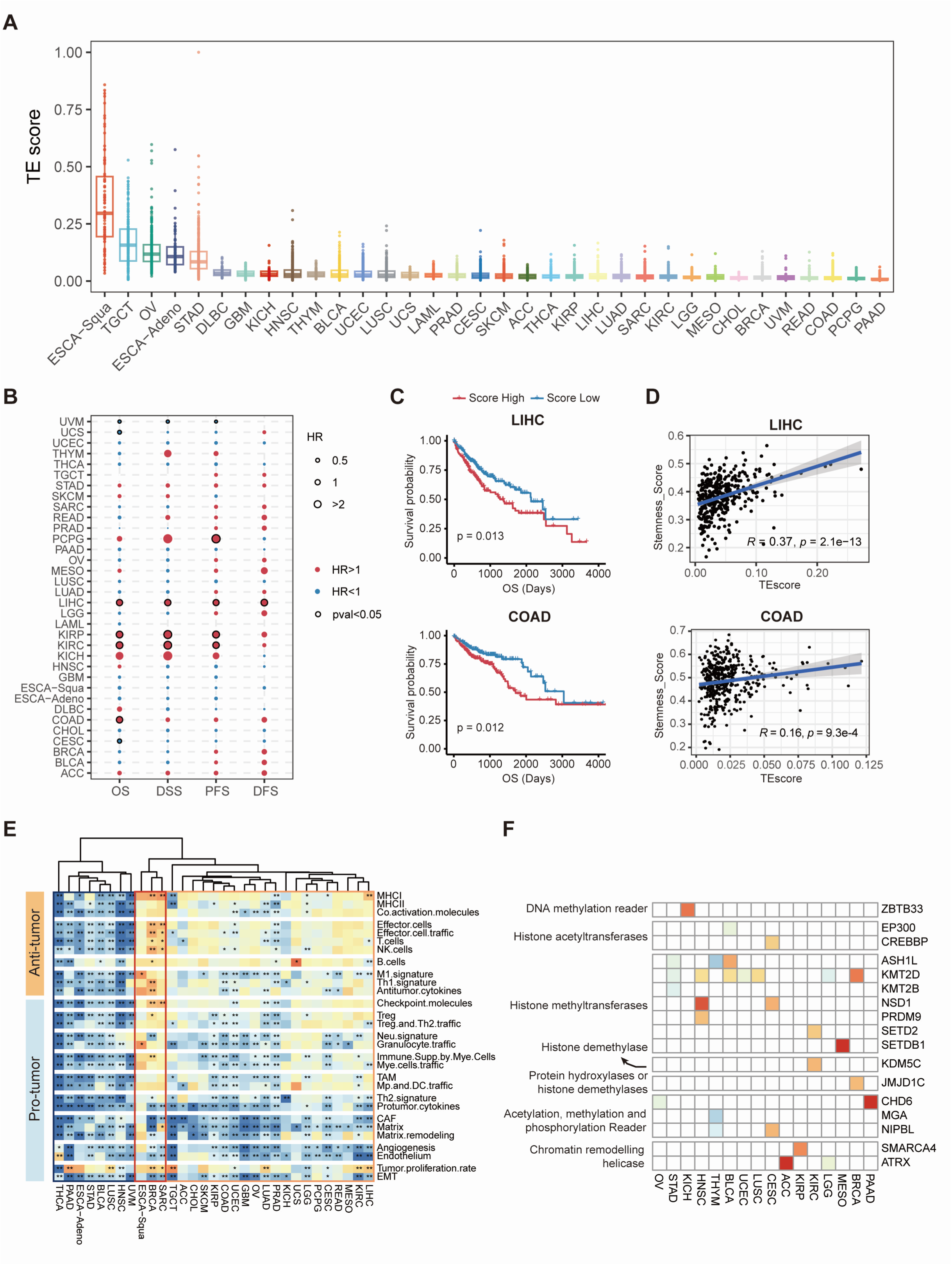
TEtrans burden (TEscore) is heterogeneously associated with prognosis and immune microenvironment across tumor types. **A.** The distribution of TEscore, reflecting the transcriptional load of TEtrans, across TCGA pan-cancer samples. **B.** TEscore is significantly associated with survival in multiple cancer types. Hazard ratios (HR) and p-values were calculated using univariate Cox regression. **C.** Kaplan-Meier curves show that high TEscore is significantly associated with poorer prognosis in liver cancer and colon cancer, with p-values determined by log-rank test. **D.** TEscore is significantly correlated with tumor stemness in liver and colon cancers. **E.** Hierarchical clustering of the correlation between TEscore and 29 tumor microenvironment pathways across TCGA cancer types. TEscore is associated with anti-tumor immune activation in esophageal squamous cell carcinoma and breast cancer, whereas with immune exhaustion in cancers such as esophageal adenocarcinoma, bladder cancer, and gastric cancer. **F.** Correlation of TEscore with epigenetic enzyme genes in various cancer types. The heatmap shows the TEscore fold change between samples with a gene mutation and its wild-type counterparts.

Correlation between TEscore and other tumor molecular features were further analyzed. We found positive associations of TEscore with tumor stemness in 16 tumor types **(Fig. 3D, Fig. S4A)**. Genomic features including aneuploidy, fraction altered, intratumor heterogeneity and homologous recombination deficiency were all more prominent in tumors with higher TEscore, indicating the chromosome instability (CIN) subtype of these tumors **(Fig. S4B)**. Tumors with mutations on some driver genes such as TP53 and epigenetic regulators such as KMT2D and SETD2, were observed with higher TEscore than their wild-type counterparts **(Fig. 3F, Fig. S4D)**. The epigenetic deregulation and TEs derepression brought by these mutations may denote the origin of TEtrans.

### TEtrans were derived from LTR elements and dual-regulated by stemness and inflammation-related transcription factors

Given our observation of inter-tumor heterogeneity of TEtrans in the transcript spectrum as well as the prognostic and immune associations, we hypothesized that tumor-type-specific expression regulation might be responsible for heterogeneous transcript with distinct functions. To explore this, we focused on the characteristics of TEtrans promoter regions in this section.

Analyzing the TE class distribution around the TSS regions (±100 bp) of TEtrans, we found that nearly half of the TEtrans were originated from LTR and Retroposon classes, with this proportion notably elevated in tumor-enriched TEtrans **(Fig. 4A)**. Further examination of a broad promoter region (±2 kb from TSS) in tumor-enriched TEtrans revealed an enrichment of ERV families, such as ERV1 and ERVK, and the LINE1 family across all examined tumor types while the SVA family from the Retroposon class particularly in ESCA, STAD, HNSC and LUSC **(Fig. 4B)**.

**Figure 4:**
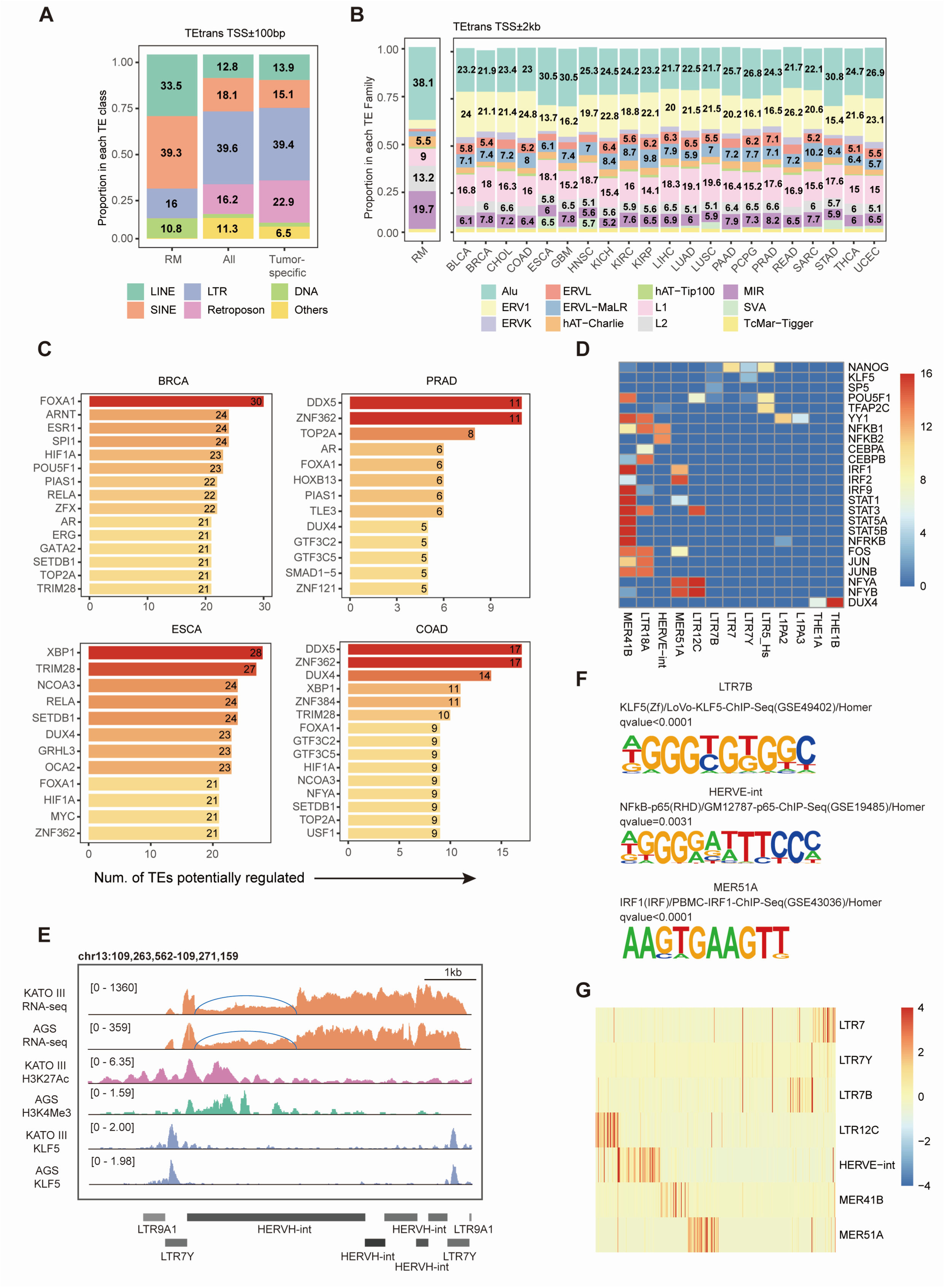
TEtrans were preferentially initiated from LTR elements and dually regulated by stemness and inflammation-related transcription factors in an element-specific manner. **A.** Distribution of TEclasses within 100bp near the TSS of TEtrans. **B.** Distribution of TE families in the promoter region (TSS ± 2kb) of TEtrans across different cancer types. **C.** Top 10 transcription factors most frequently regulating TE subfamilies across cancer types, demonstrating a cancer-type specific enrichment of vital oncogenic factors. **D.** Heatmap displaying transcription factor enrichment in the TEtrans promoter region across pan-cancer types. The x-axis represents TE subfamilies, and the y-axis represents transcription factors. Colors indicate the number of cancer types in which a specific transcription factor is enriched for a given TE subfamily, reflecting recurrent TE-transcription factor regulatory patterns across cancers. **E.** RNA-seq and ChIP-seq validation in gastric cancer cell lines showing transcriptional activity at the LTR7Y element and KLF5 binding peak enrichment. **F.** Homer motif analysis for TEtrans promoter regions, focusing on LTR7B, HERVE-int, and MER51A. **G.** The inflammatory transcription factor-regulated TEtrans and stemness factor-regulated LTR7-derived TEtrans are distributed in distinct gastric cancer samples.

TEs have been reported containing abundant docker sites for transcription factors. we further assessed how the expression of these TEtrans is regulated by measuring the propensity of transcription factors binding to TE subfamilies located within TEtrans promoter regions. Utilizing the ReMap atlas, which compiles 1210 transcription factor (TF) cistromes across 737 cell lines and tissues, we identified top transcription factors impacting the widest range of TE subfamilies for each tumor type **(Fig. 4C)**. Vital oncogenic factors are extensively enriched in a tumor-specific manner, such as ESR1 in BRCA, AR in PRAD and FOXA1 in gastrointestinal tract and prostate tumor, consistent with the tumor specificity of TEtrans expression pattern. Conversely, we observed that for particular TE subfamilies, the enrichment of TF binding regions demonstrate high similarities among different tumor types **(Fig. 4D)**. For instance, LTR7-like elements (LTR7/LTR7Y/LTR7B/LTR7C) tend to be bound by TFs associated with pluripotency and lineage differentiation, including NANOG, SOX-related families, KLF families and HOX families, in gastrointestinal tumors, lung adenocarcinoma and bladder cancer. In contrast, cistromes of inflammatory TFs such as NFKB1, NFKB2, STATs and IRFs were concurrently enriched in HERVE-int, MER41B and MER51A in a pan-cancer way **(Fig. 4D)**. We also validated these TF binding sites in LTR7-like, HERVE and MER41B elements via motif analysis and ChIP-seq data **(Fig. 4E-F)**. Such results suggest that both stem-like reprogramming and inflammatory pathways can be common mechanisms regulating TE reactivation and the consequent TEtrans expression across tumor types. Notably, TE subfamilies associated with these two common mechanisms are expressed exclusively in gastric cancer tissues **(Fig. 4G)**, further suggesting the heterogeneous regulation over TEtrans which may harbor functional implications.

### tsTE1 was a potent oncogene via strengthening the function of TOP1

We noticed that HERVH subfamily including its LTRs (LTR7/LTR7Y/LTR7B) accounted for the initiation of nearly a quarter of TEtrans in colorectal cancer **(Fig. 5A)**. Among them, a transcript originating from LTR7Y and undergoing splicing within HERVH-int demonstrated highest expression level and frequency **(Fig. 5B)**. This TEtrans, named tsTE1, was negatively correlated with both overall and progression-free survival in colorectal cancer and also abundantly expressed in multiple tumor types in a tumor-specific manner, underlining its tumor-promoting potential to be further explored **(Fig. 5C-D, Fig. S5F)**.

**Figure 5:**
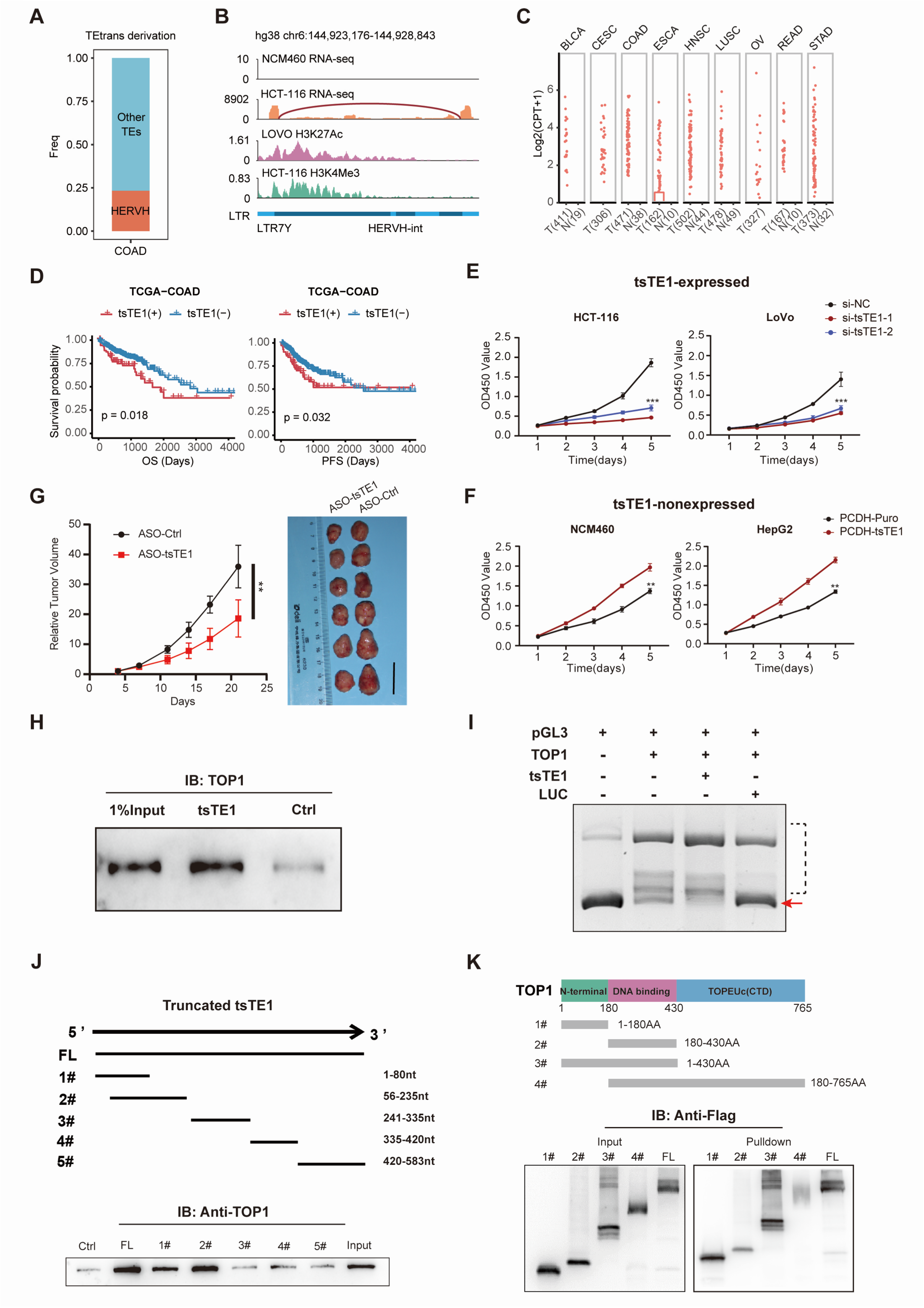
A typical TEtrans, tsTE1, was a potent oncogene via strengthening the function of topoisomerase I relaxing DNA supercoils. **A.** The HERVH subfamily initiates nearly a quarter of TEtrans events in colorectal cancer. **B.** Schematic representation of tsTE1 expression in the tsTE1-positive HCT-116 cell line and the tsTE1-negative NCM460 cell line. **C.** Boxplot showing the tumor-specific overexpression of tsTE1 in multiple cancer types. **D.** Kaplan-Meier curve showing that tsTE1 expression in the TCGA colorectal cancer cohort is associated with poor overall survival (OS) and progression-free survival (PFS). **E.** siRNA-mediated knockdown of tsTE1 in HCT-116 and LoVo cells results in decreased proliferation, as shown in proliferation assays. **F.** Overexpression of tsTE1 in the tsTE1-negative NCM460 and HepG2 cell lines significant boosts their proliferation. **G.** ASO-mediated inhibition of tsTE1 expression significantly reduces tumor volume in mouse xenograft models, compared to the control group (scale bar = 2 cm). **H.** Immunoblotting confirming the interaction between tsTE1 and TOP1 protein. LUC RNA was employed as control. **I.** Pre-incubation of TOP1 protein with either tsTE1 or control RNA followed by testing its DNA relaxing activity over the PGL3 plasmid. Results are shown by agarose gel electrophoresis. Red arrows indicate supercoiled DNA, and the black dashed box highlights the relaxed and nicked DNA bands. **J.** Immunoblotting interrogating the interaction between full-length and truncated forms of tsTE1 with TOP1 protein. **K.** Interaction of full-length and truncated TOP1 proteins with tsTE1. After transient expression of full-length and truncated TOP1 proteins, the tsTE1 RNA pulldown assay was performed, followed by immunoblotting to detect the Flag-tagged TOP1 proteins. The left panel shows input immunoblot results, confirming similar abundance of full-length and truncated TOP1 in the cell lysates used for RNA pulldown. The right panel shows the tsTE1 pulldown products, with the highest enrichment observed for truncated forms 1# and 3# relative to input.

Bioinformatics analysis predicts tsTE1 as a non-coding RNA. Having assessed tsTE1 expression across normal and cancer cell lines **(Fig. S5A)**, we selected tsTE1-expressing HCT-116 and LoVo cells, and non-expressing NCM460 and HepG2 cells for further experiment. We cloned and profiled full-length sequence of tsTE1 from RACE analysis **(Fig.S5B-C)**, and then validated the expression of 583nt-length transcript via Northern Blot in colorectal cancer cell lines and in three pairs of colorectal cancer tissue samples **(Fig. S5D-E)**. These results confirmed the validity as well as the tumor-specific feature of tsTE1 transcript.

To explore the potential oncogenic role of tsTE1, we firstly examined its effect on cell proliferation in vitro. CCK-8 assays showed that silencing tsTE1 significantly inhibited proliferation in tsTE1-expressing cells including HCT-116 and LoVo cells **(Fig. 5E)** whereas the stable overexpression of tsTE1 in non-expressing cells NCM460 and HepG2 cell lines enhanced their proliferation **(Fig. 5F)**. The EdU incorporation assays demonstrates that silencing tsTE1 significantly inhibited DNA synthesis in HCT-116 cells **(Fig. S6A-B)**. Further in vivo, we established the HCT-116 subcutaneous xenografts in which LNA-modified ASOs silenced tsTE1 and significantly inhibited tumor growth **(Fig. 5G)**. Both in vitro and in vivo experiments demonstrated that tsTE1 can markedly boost the proliferative phenotype of colorectal cancer cells.

For mechanism exploring, we initially performed RNA pulldown assay to identify the proteins possibly binding to tsTE1. Notably, a distinct protein band at approximately 100 kD was more significant for tsTE1 than its antisense transcript **(Fig. S6C)**. Subsequent mass spectrometry and immunoblotting revealed that DNA topoisomerase I (TOP1), an essential enzyme responsible for the relaxation of supercoils during cell cycle ^31^, could specifically bind to tsTE1 **(Fig. 5H)**. We conducted an in vitro plasmid relaxation assay to assess if the tsTE1-TOP1 interaction alters TOP1’s function **(Methods)**. The results showed that tsTE1 pre-incubation stimulated TOP1 relaxation above its basal activity, suggesting that tsTE1 may promote tumor proliferation by boosting TOP1’s supercoil relaxation activity **(Fig. 5I)**.

To investigate how tsTE1 can affect TOP1 activity, we constructed a series of truncated variants of tsTE1 based on its secondary structure and performed RNA pulldown assays separately **(Fig. S6D-E)**. Of note, 56-235nt of tsTE1 was the critical region for the interaction with TOP1, corresponding to the first exon of the transcript **(Fig. 5J)**. In parallel, we created TOP1 truncated variants to pinpoint the primary interacting domains **(Fig. 5K)**. Pulldown assays with these deletions revealed that removing the N-terminal domain significantly attenuated the co-immunoprecipitation of TOP1 with tsTE1, whereas deletion of the C-terminal enzymatic domain had minimal impact on the interaction (**Fig. 5K**). These results indicated that the N-terminal domain of TOP1 facilitated the interaction with tsTE1, rather than the conserved enzymatic domain typically found in eukaryotes.

In summary, we identified a HERVH-derived TEtrans, tsTE1, promoting tumor proliferation through the interaction with the N-terminal domain of TOP1 enhancing its activity of DNA supercoil relaxation. These experimental evidence stressed that TEtrans may provide a novel source of oncogenes contributing to tumor development.

### Part of TEtrans harbor coding ability serving as a source of tumor neoantigens

As TEtrans represent a novel form of TE-derived mature transcript with tumor-enriched expression, we speculated that these transcripts may also encode tumor neoantigens potentiating anti-tumor immunity. We scanned the open reading frames (ORF) of TEtrans and predicted their coding abilities with CPAT software. After eliminating TEtrans not expressed in CCLE tumor cells and those encoding annotated peptides, we obtained 398 tumor-enriched TEtrans candidates potentially coding 387 neopeptides, accounting for 7.2% of total TEtrans **(Fig. 6A)**. The low proportion of coding transcripts could be attributed to the destruction of most original TE ORFs during evolution. However, with the independence from protein-coding genes, these TEtrans candidates can provide unrecognized neoantigens at full-length scale in contrast to TE-chimeric transcripts, conferring their advantages for tumor mRNA vaccines.

**Figure 6:**
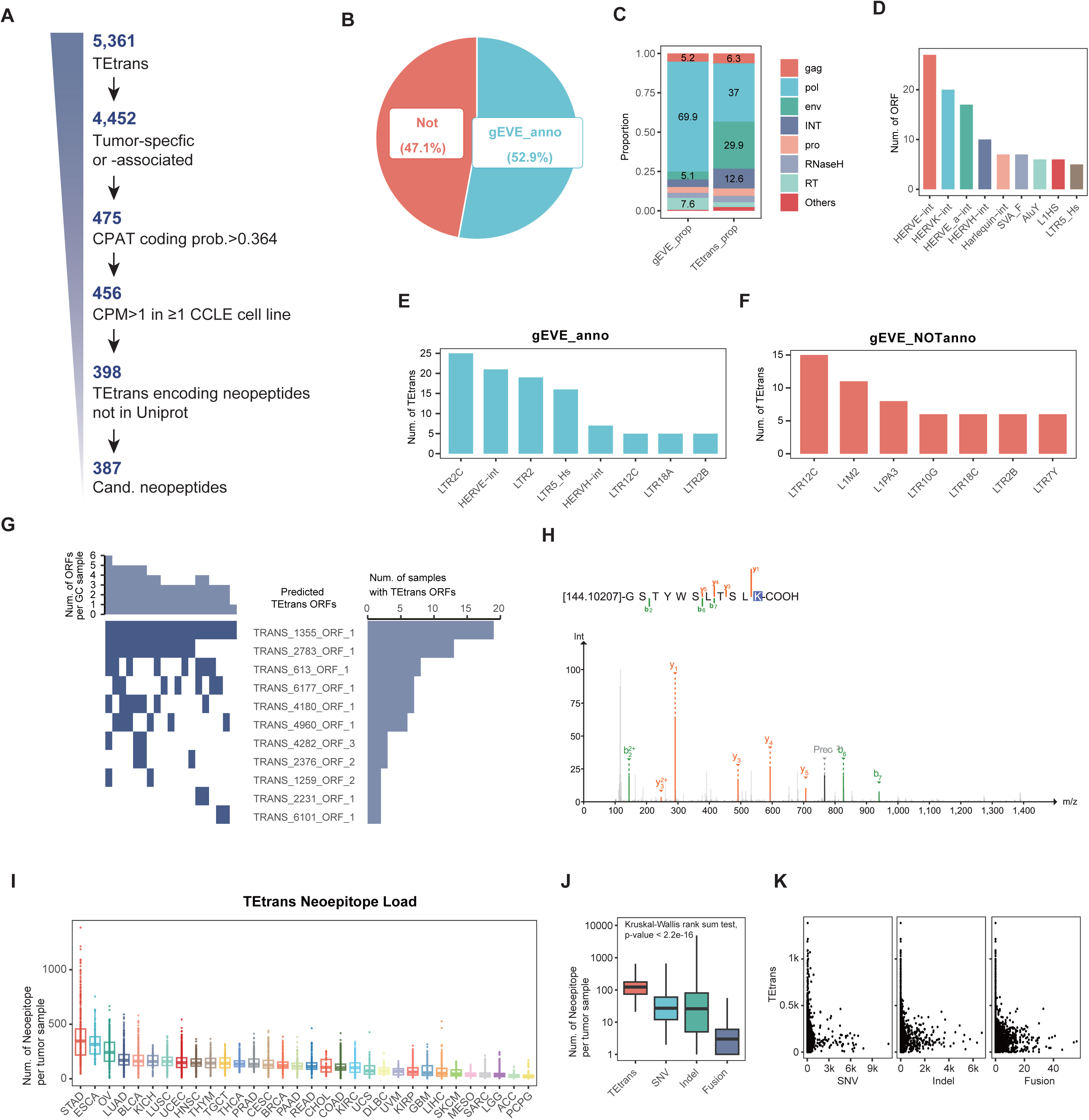
Part of TEtrans harbor coding ability serving as a source of tumor neoantigens. **A.** Identification of TEtrans with potential coding ability. A total of 398 TEtrans elements, enriched in at least one tumor type, were predicted to encode 387 novel peptides. **B.** Proportion of TEtrans coding candidates annotated to viral proteins curated in the gEVE database. **C.** Distribution of annotated viral proteins encoded by TEtrans compared to the genomic background. **D.** Viral proteins encoded by TEtrans are primarily derived from the ERV1 family. **E-F.** Distribution of the TE subfamilies across 100bp near the TSS of TEtrans coding candidates. TEtrans potentially coding viral proteins are mainly initiated from HERVE (including LTR2C/LTR2) and HERVK (including LTR5_Hs) (E), while those genrating non-viral products predominantly derived from LTR12C and LINE1 family (F). **G.** Peptides encoded by TEtrans can be recurrently detected from the proteome data of gastric cancer tissues. **H.** Mass spectrum of peptides from TRANS_4180 in a gastric cancer sample. **I.** TEtrans-derived neoepitope load across cancer types in the TCGA cohort. **J.** Tumors harbor more neoepitopes derived from TEtrans than genomic alterations (single nucleotide variations (SNV), insertions/deletions (Indels), and gene fusions) in the individual sample level. **K.** Distribution of neoepitopes derived from TEtrans, SNV, Indels, and gene fusions across tumor samples, showing that TEtrans contributed more potential neoantigens in tumors with few genomic variations.

These predicted protein-coding TEtrans were predominantly composed of two or three exons with the median ORF as 152AAs in length **(Fig. S7C-D)**. About a quarter of their ORFs spanned the TE splicing junction, indicating that the aberrant splicing of TEs is an important source of TEtrans-encoded proteins in tumors **(Fig. S7E)**. Given that endogenous retrovirus (ERV) provides the predominant genomic material for TEtrans, we presume that TEtrans may partially retain the retroviral protein coding frames. Hence, we blasted the viral ORFs curated by gEVE database^32^, containing gag, pol, and env, in these predicted coding TEtrans and obtained 52.9% of the ORF frames encoding viral proteins **(Fig. 6B)**. Compared to all annotated endogenous viral ORFs, TEtrans ORFs are highly enriched in env proteins **(Fig. 6C)**. These results suggest that although proteins coded by TEtrans may not assemble into complete viral particles, they may still exert corresponding functions encoding certain parts of viral protein. Notably, predicted coding TEtrans with viral or non-viral ORFs are derived from different TEs **(Fig. 6E-F)**. The former is mainly initiated from HERVE including the internal sequences and the flanking LTR elements (LTR2/LTR2B/LTR2C) **(Fig. 6E)**, while the latter is primarily derived from LTR12C and LINE families such as L1M2 and L1PA3 **(Fig. 6F)**. Considering that both HERVE and LTR12C were demonstrated as inflammation-associated TEs via TF enrichment analysis **(illustrated in previous section)**, we speculated that the expression of TEtrans potentially coding could be coordinated by the activation of inflammatory pathway.

To validate the presence of TEtrans-encoded proteins, we conducted mass spectrometry (MS)-based proteomic analysis on 19 tumor tissues from CPTAC early-onset STAD cohort^33^. We constructed a customized protein database comprising 106 predicted TEtrans-derived proteins, which was concatenated to the annotated human proteome (UniProt/Swiss-Prot) for MS data interrogation. Notably, peptides corresponding to 36 TEtrans-derived proteins were identified in at least one STAD tumor sample, with 11 proteins recurrently detected across multiple samples **(Fig. 6G-H)**. These findings the capability of TEtrans to encode proteins previously uncharacterized and potentially immunogenic.

To further investigate whether these TEtrans can provide neopeptides presented by MHC-I molecules, we in silico translated ORFs of TEtrans coding candidates into 9-mer peptides. These peptides predicted with high affinity to HLA-I molecules for each TCGA patient were subsequently defined as TEtrans derived neoepitopes **(Methods)**. TEtrans of STAD, ESCA and OV bear the highest burden of neoepitopes, whereas that of LIHC and SARC, in spite of a relatively high number, contain few neoepitopes **(Fig. 6I, Fig. S7A).** Such findings suggest the immunoediting of TEtrans-derived neoantigens in different tumor types. Measuring anti-tumor immunity revealed that tumors with high TEtrans neoepitope burden demonstrated increased IFN-γ repsonsiveness, high expression of immune checkpoints as well as elevated infiltration of CD8+T cells and activated DCs **(Fig. S7B)**. These results implied that the immune evasion of tumors with abundant TEtrans neoantigens was at least partially attributed to elevated immune checkpoints whereas antigen presenting machinery and T cell response could be fairly preserved. Considering that previous therapeutic vaccines were mostly developed on neoantigens derived from genomic alterations, especially mutations^34,35^, we compared the immunogenic potential of TEtrans neoepitopes with that of three common genomic variants: single nucleotide variations (SNVs), insertions and deletions (Indels), and gene fusions. TEtrans generated a higher neoepitope load per individual sample than genomic alterations meanwhile contributed more potential neoantigens in tumors with few genomic variants **(Fig. 6J-K)**, stressing the profound value of such transcripts as the complementary targets of genomic alterations for the development of TCR-T and mRNA vaccines.

### TEtrans can activate CD8+T cells capable of antigen-specific tumor lysis

For verification of these predicted coding TEtrans, we screened four tumor-enriched TEtrans which were shared among multiple tumor types and demonstrated positive associations with anti-tumor activity **(Fig. 7A, D, Fig. S8A-C)**. These four transcripts were all composed of two exons, with one of them, T4180, derived from LTR30 while the other three initiated from LTR12/LTR12C **(Fig. 7A)**. We further synthesized the corresponding mRNA by in vitro transcription and validated the protein product of these TEtrans in 293T cell line **(Fig. S8D)**.

**Figure 7:**
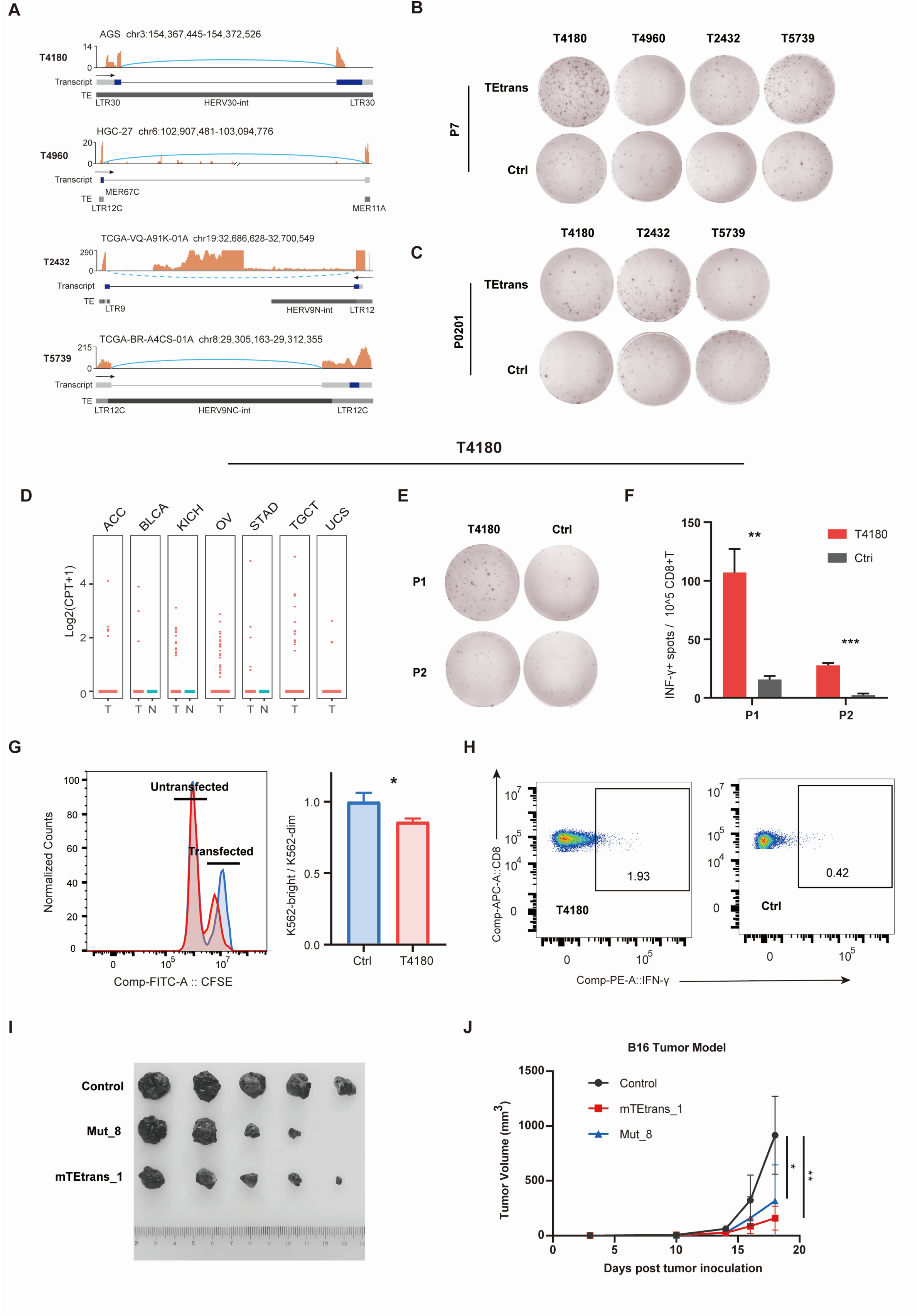
TEtrans-derived neoantigens demonstrate immunogenicity in vitro and in vivo. **A.** A schematic illustrating the transcript structures of four selected TEtrans elements and their expression in tumor tissues or cell lines. The blue rectangles represent predicted open reading frames. Positive or negative strand were denoted by solid or dashed lines respectively. Varying shades of gray represent different evolutionary ages. **B-C.** Transfection of four TEtrans mRNA candidates into Mo-DCs derived from PBMCs of two healthy individuals induces TEtrans-specific CD8+ T cells that secrete IFN-γ, indicating the presence of TEtrans-specific T precursor cells in healthy individuals. Representative ELISPOT staining results of two healthy individuals shown in panels B and C. **D.** The T4180 transcript is specifically expressed in multiple tumor types including ovarian, gastric, and testicular germ cell tumors. **E-F.** Transfection of T4180 mRNA into PBMCs of two gastric cancer patients activates CD8+ T cells correspondingly to secrete IFN-γ. Representative ELISPOT staining results (E) and spot quantification (F) are shown. **G.** T4180-induced CD8+ T cell clones specifically kill K562-HLA0201 cells expressing the corresponding antigen. The difference in the K562-bright/K562-dim ratio (T4180 vs. Ctrl) reflects the specific killing proportion. **H.** Intracellular IFN-γ flow cytometry analysis showing the reactivation of T4180-induced CD8+ T cell clones by K562 cells expressing T4180. **I-J.** Evaluation of the tumor treatment effect of mRNA vaccine based on mouse TEtrans homologous transcript (mTEtrans_1) in the B16F10 xenografts.

To validate whether these proteins can act as tumor neoantigens to activate immunity, we first constructed a mRNA immunogenicity screening platform based on human peripheral blood mononuclear cells **(Fig. S8E, Methods)**. Immunogenecity of four TEtrans was tested by directly transfecting PBMCs with pseudouridine modified mRNAs to induce the activation of T cells from two healthy volunteers, one of which harboring HLA-A02:01 genotyping. We observed that all four TEtrans mRNA could activate T cells in at least one donor to secrete IFN-γ **(Fig. 7B, Fig. S8F-G)**. As peptides derived from T4180 and T4960 were identified by MS of gastric tumor tissues, We further validated the capabilities of these two TEtrans to activate CD8+ T cells in two patients with gastric cancer. CD8+ T cells from both donors were successfully activated by peripheral B cells transfected with T4180 mRNA **(Fig. 7E-F, Fig. S8H)**. These results suggest that TEtrans are immunogenic in vitro and and some of them such as T4180 can generate epitopes presenting by multiple HLA-I genotypes. TEtrans-specific TCRs can exist in the TCR repertoire of healthy donors which could recognize TEtrans-derived antigen epitopes.

To confirm the specificity and killing efficacy of TEtrans-induced CD8+ T cell clones, we chose HLA-A*02:01, the most common HLA-I genotype in the population, for tumor killing assay **(Fig. S8I, Methods)**. Two TEtrans eliciting best T cell response in PBMCs of HLA-A*02:01 healthy donors, T4180 and T2432, were subjected to verification. Both two TEtrans-specific CD8+ T cell clones exhibited antigen-specific cell lysis and elevated IFN-γ secretion when incubated with K562 cells expressing corresponding TEtrans compared to cells not transfected, supportive of the immunogenicity of TEtrans **(Fig. 7G-H, Fig. S8J-K)**.

The primate-specific feature of TEtrans can hinder the direct validation of their immunogenic potential in mouse models. Nevertheless, we attempted to identify homologous transcript mimicking TEtrans features in mouse tumor cells. A transcript initiated from and exclusively spliced among TEs, namely mTEtrans_1, was highly expressed in B16F10 mouse melanoma cell line. The mRNA vaccine was designed based on mTEtrans_1 and its tumor-suppressing effect was subsequently measured in C57 syngeneic model with B16F10 tumor cells **(Fig. S9A)**. Strikingly, the mTEtrans_1 vaccine demonstrated tumor control comparable to that encode combinations of eight somatic mutations, inducing drastic tumor regression after twice subcutaneous injection **(Fig. 7I-J)**. An elevated proportion of IFN-γ+ splenic CD8+ T cells was also detected after mTEtrans_1 reactivation compared with control **(Fig. S9B)**. Such positive tumor response of mTEtrans_1 vaccine further implied the promising value of TEtrans as targets for mRNA-based therapeutic vaccine.

## Discussion

TEs, constituting nearly half of the human genome, are reactivated in tumors and profoundly contribute to transcriptomic heterogeneity. While extensive studies have focused on TE-gene chimeric transcripts, the landscape of gene-independent TE-derived transcripts remains underexplored. Attempts to systematic characterize such transcripts have been hampered by technical limitations, including excessive filtering of repetitive sequences and ambiguous quantification of TE-derived reads. Here, leveraging our splicing junction-centric analytical framework (ASJA), we characterized a tumor-enriched gene-independent TE-derived transcript form which are autonomously transcribed and spliced from TEs across 32 tumor types resulting 5,361 transcripts. These transcripts, termed TEtrans, predominantly unannotated (>90%) and enriched in intergenic regions, represent a novel layer of TE-mediated transcriptomic innovation. Their expression are epigenetically activated and dually regulated by stemness/inflammation-related transcription factors, suggesting context-dependent functional roles. We further illustrated how TEtrans can act as a “double-edged sword” with both oncogenic and tumor-suppressive capacities through dissection of the tumor-promoting mechanisms exemplified by HERVH-derived tsTE1 and validation of the immunogenicity of TEtrans-encoded peptides, respectively. These findings not only extend the functional landscape of TEs but also illuminate novel therapeutic avenues — either by exploiting their immunogenic properties for vaccine development or targeting oncogenic isoforms through precision interventions.

TEs have been well-recognized as contributors of alternative splicing events within gene regions ^25,36^. Notably, TE splicing events in TEtrans predominantly occur in intergenic regions, extending the scope of TE involvement beyond gene-specific alternative splicing. In contrast to previously described TE-exon chimeric splicing (JETs) which involve mostly intronic TEs with junction breakpoints distributed disperse ^16^, TEtrans are mostly spliced from primate-specific and evolutionarily younger TEs, especially SVA and ERV families, with highly conserved splice site motifs. Notably, some TEtrans splice sites align with those identified in intact retroviral RNAs, implying that TEtrans may arise from cryptic splicing sites of newly integrated TEs due to low mutation accumulation and incomplete epigenetic silencing ^37,38^. These conserved junction breakpoints further suggests the involvement of tumor-specific post-transcriptional machinery^39^, warranting further investigation through RNA-binding protein profiling (e.g., CLIP-seq) and functional studies.

Documented as cis-regulatory elements (cCREs), TEs have been reported to function as promoters or enhancers providing TF binding sites to regulate downstream targets^40^. To investigate the regulatory basis of TEtrans expression, we performed TF motif enrichment analysis on TEs within TEtrans promoter regions. These regions were enriched for binding sites of lineage-specific oncogenic TFs in chromatin-accessible TEs, aligning with the observed tissue-specific expression patterns of TEtrans. These findings suggest that, while TEtrans represent a novel regulatory innovation of TEs, their expression is predominantly governed by a canonical mechanism wherein epigenetically reprogrammed TEs are co-opted by lineage-specific TFs—a phenomenon previously reported in fetal development^41^ and multiple cancer types^42,43^. Intriguingly, TEtrans regulated by stemness-associated TFs (e.g., NANOG, KLF5) and inflammation-related TFs (e.g., STAT3) originate from distinct TE families and are exclusively expressed in tumors. This functional dichotomy parallels the established roles of other TE-derived transcripts in tumor promotion and immune modulation^12,44^, leading us to hypothesize that stemness- and inflammation-associated TEtrans may play divergent roles in tumorigenesis. Such functional heterogeneity underscores the need for mechanistic studies to dissect their context-dependent roles in tumor biology.

The HERVH and its LTRs are known to participate in the maintenance of pluripotency in stem cells ^45–48^. In colon cancer—a tumor type tightly linked to stemness—we found that HERVH initiates ∼25% of TEtrans, far exceeding other TEs. While HERVH activation has been implicated in colon cancer progression^22,49,50^, the specific transcript mediating its tumor-promoting effects remains elusive. Through our junction analysis, we identified multiple HERVH-derived TEtrans, including tsTE1, a highly expressed and recurrent unannotated transcript. Functional studies revealed that tsTE1 promotes colorectal cancer proliferation by interacting with TOP1 and enhancing its DNA supercoil relaxation activity. Although emerging evidence has suggested the role of TOP1 as an RNA-binding protein, its RNA-mediated regulation of DNA topology is poorly characterized. A recent study reported that TOP1 binding to mRNA via its DNA-binding core domain inhibits DNA relaxation during transcription^51^. In contrast, tsTE1 interacts with the N-terminal domain of TOP1, a region previously associated with stimulation by proteins like nucleolin^52^ and subunits of preinitiation complex (e.g. RNAPII, TFIID)^53,54^, to amplify DNA relaxation efficiency. This distinct mechanism suggests that TEtrans may exploit TOP1’s modular domains to fine-tune genomic topology in a context-dependent manner. Given the abundance of HERVH-derived TEtrans, we propose that similar interactions could underlie broader oncogenic functions of this TE family. Together, these findings establish tsTE1 as a critical effector of HERVH-driven tumorigenesis and position TEtrans as a reservoir of unexplored TE-derived oncogenes.

Beyond their tumor-promoting roles, TEtrans represents a novel source of "non-self" proteins providing tumor neoantigens considering their tumor-specific expression and recurrence in the population. Approximately 5% of TEtrans which predominantly derived from ERV1 and L1 families are predicted to encode proteins, with ORFs verified by MS data of gastric cancer. Despite that the evolutionary pressure tend to disrupt TE ORFs limiting their coding potential, TEtrans are potent to provide more unrecognized neoepitopes for their gene-independent feature. This potential was supported by our observation that TEtrans surpass genomic alterations in neoepitope yield per tumor sample, especially in tumors with a low mutational load, establishing them as a valuable source of tumor neoantigens. Functional studies further suggest that proteins encoded by TEtrans can elicit antigen-specific immune responses. We developed an mRNA-based pipeline that simulates the endogenous antigen expression, processing and presentation^55^, contrasting with traditional validation methods directly loading APCs with peptide epitopes^14,16,56,57^. Our results demonstrated that TEtrans-derived antigens activate CD8+ T cells in both healthy donors and tumor patients, with induced CD8+ T cell clones specifically lysing TEtrans-expressing tumor cells. Despite the primate specificity of TEtrans, we identified murine transcripts in B16F10 cell lines mimicking TEtrans features. One such transcript exhibited in vivo antitumor efficacy comparable to hotspot mutations, providing cross-species immunogenicity validation. Reprogrammed TEs are emerging as a critical source of tumor neoantigens. Beyond TE-exon chimeras widely studied, our work introduces a type of autonomous TE-derived and spliced transcripts as antigen-generating entities independent of host gene frameworks—expanding the potential applications of TEs for therapeutic vaccination and TCR-based cell therapies.

In summary, our study characterized TEtrans, a prevalent class of TE-derived transcripts independent of gene context in tumors, investigated their conserved splicing patterns and epigenetic regulation, and finally uncovered their dual functional roles as reservoir of both oncogenes and tumor neoantigens. As transcripts autonomously transcribed and spliced from TEs, TEtrans are mostly derived from primate-specific TEs and exhibit tumor-specific expression, decoupling TE activity from host gene frameworks and representing a novel layer of adaptive tumor evolution. Functional findings further underscore the potential of TEtrans as promising targets for designing precision interventions, therapeutic vaccinations and T cell-based modalities across a wide range of cancer types.

## Materials and Methods

### Data acquisition

We downloaded the original bam sequencing files of 33 tumor types and corresponding adjacent tissue samples from the GDC (https://gdc.cancer.gov). Raw RNA sequencing data of 30 normal tissues covering 17,382 samples were acquired from the Genotype-Tissue Expression (GTEx) portal website (https://www.gtexportal.org/). Raw RNA-seq data of 1,046 cancer cell lines were retrieved from the CCLE website (https://portals.broadinstitute.org/ccle).

### The identification of TEtrans

Splicing events in TCGA, GTEx, and CCLE samples were identified using the ASJA program^26^. The analysis proceeded as follows: Raw FASTQ sequencing files were first assessed for quality using FastQC. The reads were then aligned to the hg38 reference genome using the STAR aligner (two-round alignment), generating BAM files. These BAM files were merged with StringTie according to the GENCODE v29 annotation. The coverage of linear splice sites was calculated based on the minimum exon coverage at both splice site ends and normalized to the coverage of all annotated splice sites, yielding the CPT-normalized splice site expression.

Next, we extracted the donor and acceptor sites of splicing events, constructed BED files, and used the bedtools intersect command to map the genomic positions to the human hg38 TE annotation. A splice site was classified as a TE splicing event if at least one end of the splice site overlapped with a TE. The TE annotation file was downloaded from the UCSC Genome Browser and annotated using RepeatMasker software. This file includes information on the genomic location, family, class, and age of the TEs. TEs were classified into broad categories based on sequence similarity: Long Interspersed Elements (LINEs), Short Interspersed Elements (SINEs), Long Terminal Repeats (LTRs), DNA elements, and retroposons, with further subdivisions into families and subfamilies. For this study, TEs of the same family with distinct differentiation ages were classified as subfamilies to include age-related information. To identify TEtrans, transcripts corresponding to TE splicing events were filtered according to the following criteria: (1) Both the upstream and downstream 100 bp of the transcription start site (TSS) were located within a TE; (2) All splice sites were positioned within the TE region; and (3) More than 50% of the transcript content consisted of TE sequences.

### Evolutionary age of TEs

We referred to the method described by Pierre-Emmanuel Bonte et al.^21^ to calculate the evolutionary age of TEs using the formula: Divergence/(2.2*10^^-9^). Divergence represents the sequence difference between a TE and its closely related element.

### Characterizing TEtrans

#### MEME motif enrichment analysis

The FASTA sequences of TEtrans were extracted from the reference transcriptome using the gffread tool. Subsequently, TEtrans sequences were analyzed for motif enrichment using the STREME tool from the MEME Suite^58^. Following the significant enrichment of the AAUAAA motif, a well-known polyadenylation signal, we extracted sequences upstream of the transcription termination site of TEtrans, extending 40 bp or 150 bp, respectively. To assess the enrichment of the polyadenylation signal, we used the SEA tool with a scrambled base sequence as the background. Sequences containing the AAUAAA motif were identified. We randomly selected equal numbers of mRNAs and lncRNAs, extracting sequences from the same regions of these transcripts for comparison.

#### FANTOM5 CAGE peak analysis

Cap Analysis of Gene Expression sequencing (CAGE-seq) provides single-base resolution of transcription start sites (TSS) and allows quantification of transcriptional activity at each TSS. The FANTOM5 (Functional Annotation of the Mammalian Genome 5) project^59^ has cataloged over 1,800 human tissues, primary cells, and tumor cell lines to comprehensively map transcriptional activity across diverse cell types and tissues using CAGE technology. To further validate the authenticity of the TEtrans transcript, we obtained the FANTOM5 TSS location data (https://fantom.gsc.riken.jp) and identify intersections between regions around TEtrans TSS (defined as TSS±500 bp) and the FANTOM5 TSS peaks. TEtrans regions with such intersections were considered CAGE-associated, thereby confirming the transcription initiation of TEtrans.

#### ATAC-seq and ChIP-seq data analysis

ATAC-seq (Assay for Transposase-Accessible Chromatin with high-throughput sequencing) leverages the preferential insertion of transposase Tn5 into open chromatin regions to assess the accessibility of chromatin regions for transcription factors and other regulatory proteins. To evaluate chromatin accessibility around the TSS of TEtrans, we retrieved chromatin accessibility data^60^ for 655 samples across 22 tumor types from the Genomic Data Commons (GDC)(https://gdc.cancer.gov/about-data/publications/ATACseq-AWG/). Additionally, we performed H3K27Ac ChIP-seq on the HUH7 cell line as a marker of active chromatin. We then analyzed the ATAC-seq data from TCGA samples and the H3K27Ac ChIP-seq data from HUH7 cells, focusing on the TSS regions of TEtrans, defined as ±2kb from the transcription start site. Signal intensity was quantified by calculating the signal enrichment scores for these regions using the computeMatrix function from the deeptools software. The resulting data was visualized by plotHeatmap.

#### Third-generation validation

Third-generation full-length RNA sequencing directly sequences RNA molecules, offering longer read lengths that enable more accurate identification and differentiation of TE-derived transcripts. To compare the transcriptomic profiles, the HUH7 cell line was subjected to both standard next-generation transcriptome sequencing and third-generation full-length RNA sequencing. For the third-generation sequencing data, long reads were aligned to the human reference genome (hg38) using the minimap2 command within the FLAIR program. Linear splice sites were then extracted from the alignment file using the 2passtools tool. RNA-seq data for both cell lines were analyzed following the ASJA protocol.

### Conservation analysis of TEtrans splicing positions and motifs

The consensus sequence represents the typical characteristics of a transposable element (TE) and serves as a critical reference for systematically analyzing copies of the same TE. We retrieved TE HMM models from the Dfam website (Dfam 3.7 Annotations) and then mapped splicing position on each copy to corresponding location on the consensus sequence. Locations frequently distributed were considered as conservative splicing events. For motif identification of splicing sites, we categorized each TEtrans splicing event by TE families and splicing modes. To characterize the splice sites, we extracted 9 bases from −3 to +6 base pairs relative to the donor site, defining this region as the 5’ splice site (5ss). Similarly, we extracted 43 bases from −40 to +3 base pairs at the acceptor site, defining it as the 3’ splice site (3ss). Sequence conservation of splicing motifs were visualized with ggseqlogo R package across the different TE trans splicing events.

### Validating TEtrans conserved splicing motifs

To verify the splicing efficiency of the TEtrans splice sites, we selected conserved 5ss and 3ss motifs, paired them based on the TE splicing pattern, synthesized artificial introns, and inserted them into the EGFP coding sequence. Splicing efficiency was assessed in 293T cells by comparing the proportion of amplification products across splice sites to total EGFP products, analyzed qualitatively via agarose gel electrophoresis and quantitatively by qRT-PCR.

### Assessment of prognostic and molecular associations with high TEtrans load

We firstly developed TEscore to evaluate the overall TEtrans load considering both expression level and frequency. The expression value of each TEtrans was standardized at the pan-cancer level using the z-score normalization. Subsequently, z-scores of all TEtrans were summed within each sample. We performed the max-min normalization across samples to ensure that the final TEscores ranged from 0 to 1, with higher TEscore values indicating a greater transcriptional burden of TEtrans.

TEscore was divided into two groups based on the median value. The prognostic significance was analyzed by univariate Cox regression. Survival curves were generated using Kaplan-Meier analysis, with significance determined by the Log-rank test. The association between TEscore and stemness score was assessed in each TCGA tumor type using Spearman correlation analysis. Aneuploidy scores, genomic copy number variation (CNV), intratumor heterogeneity, and homologous recombination deficiency were compared between the high and low TEscore groups subjected to Wilcoxon nonparametric tests.

Relationships between tumor TEtrans load with mutation characteristics were evaluated in 618 cancer driver genes obtained from IntOGen (https://www.intogen.org/download) and 212 epigenetic enzyme genes compiled by a previous study^61^. For each key mutant gene, we calculated the fold change in TEscores between samples with and without gene mutations and statistically compared them with Wilcoxon nonparametric test.

### TF enrichment analysis for TEs potentially controlling TEtrans expression

TEtrans represents a novel TE-autonomous transcript form, the expression of which was potentially attributed to cis-regulatory co-option of TEs in tumors. To investigate the expression control of TEtrans, we conducted an enrichment analysis of transcription factors over TEs located in the promoter regions of TEtrans (TSS±2kb) integrating peak sets of each TF and ATAC-seq data of each tumor type, referring to methods from a previous work^42^. Across 17 tumor types with available ATAC-seq data from TCGA (including BLCA, BRCA, CHOL, COAD, ESCA, GBM, HNSC, KIRC, KIRP, LIHC, LUAD, LUSC, PCPG, PRAD, STAD, THCA, and UCEC), we utilized BEDtools intersect command to identify TEs that intersected with both the promoter regions and ATAC-seq peaks specific to each tumor type. All TEs annotated in Repeatmasker aggreeing with such standards were extracted as the background. The Bioconductor package LOLA^62^ was then employed to perform TF enrichment analysis between query TEs and background. TF binding sites were obtained from the ReMap database^63^ (remap2020 - http://remap.univ-amu.fr/), which catalogues ChIP-seq peaks for 1,135 TFs. The command runLOLA (query, univ, regionDB, cores=1) was used to determine the odds ratio (OR) and significance of each TF enrichment for each TE relative to the background.

### TF motif enrichment analysis

The TF motif enrichment was performed using the Homer software^64^ with the command findMotifsGenome.pl TE.bed -bg background.bed -size 200. The enrichment results for known transcription factor motifs in TEs were obtained from the generated knownResults.txt file.

### Prediction of protein-coding ability for tumor-enriched TEtrans

We extracted the sequences of tumor-enriched TEtrans using the gffread software^65^ and predicted their protein-coding ability and open reading frames (ORFs) of TEtrans with the CPAT software^66^. We performed in silico translations for TEtrans sequences and ORFs with a start codon of ATG and a length greater than 30 bp were identified. Following the recommendations from CPAT, we selected ORFs with a coding ability score greater than 0.364 as potential protein-coding ORFs. TEtrans containing such ORFs were defined as potential protein-coding TEtrans for subsequent analysis and experimental verification.

### Blasting viral proteins in TEtrans ORFs

We retrived viral ORFs from the gEVE database curating 31,292 non-overlapping ORFs of viral proteins identified in the human genome. We downloaded the fasta file containing the human genome viral protein sequences and aligned TEtrans-encoded ORFs against the viral protein database using the blastp software. An ORF was defined as encoding a viral protein if it had an identity percentage (pct_identity) >85% and a mismatch count ≤2 during alignment.

### MS-based validation of TEtrans-derived peptide

Paired transcriptome and proteome data of 19 tumor tissues derived from patients with early-onset STAD were retrieved from PDC portal (https://pdc.cancer.gov/pdc/). RNA sequencing data were subjected to ASJA with TEtrans expressed in at least one tumor sample were included to construct protein database. The database containing 106 predicted proteins translated from TEtrans were concatenated to UniProt/Swiss-Prot proteins for MS-based peptide identification. The mgf files were searched against the unified protein database using msfragger software. The parameters for peptide identification were adopted according to the work of Dong-Gi Mun et al.^33^. Briefly, the true precursor mass tolerance was set to 10ppm and semi-enzymatic termini was allowed. Oxidation on methionine and loss of amonia on glutamine and asparagine were included as variable modifications whereas iTRAQ on lysine and N-terminalof peptide as well as carbamidomethylation on cysteine were added as static modifications. The MS spectrum was visualized by PDV software (V2.1.0).

### Prediction of TEtrans neoepitopes

We first generated TEtrans-derived peptide database (TEtrans_db) through digesting TEtrans-encoded ORFs into overlapping 9-mer peptides as the optimal MHC class I epitopes are typically 9 residues in length. Peptides absent from UniProt/Swiss-Prot were retained for subsequent MHC affinity prediction. We obtained the four-digit HLA class I genotyping data for 8,915 TCGA tumor samples from the study by Thorsson et al.. Then peptide-MHC affinity was predicted separately for each patient based on their corresponding HLA alleles. We employed NetMHCpan (v4.1) software to predict MHC affinity of TEtrans_db for each patient based on their corresponding HLA alleles. Peptides predicted with strong binding and affinity below 500 nM were considered as neoepitopes of TEtrans. Estimation of tumor neoantigens derived from mutations, indels and fusions for TCGA tumor samples were obtained from the studies by Thorsson et al.^67^ and Yang et al.^56^.

### Experiemental Procedures

#### Cell Culture

HEK293T cell line was obtained from the ATCC. HuH7, HepG2, HCT116, LoVo and NCM460 cells were purchased from the Shanghai Cell Bank Type Culture Collection (Shanghai, Chinese Academy of Sciences, China). The authentication of all cell lines were ensured by short tandem repeat profiling. Mycoplasma contamination was regularly examined using qRT-PCR. Cells were maintained in appropriate medium supplemented with 10% FBS, 100 mg/mL penicillin and 100 U/mL streptomycin.

#### Rapid amplification of cDNA ends (RACE)

We applied 5’- and 3’-RACE assays to amplify the full length cDNA sequence of tsTE1 from the RNA of HCT116 cells using the SMARTer™ RACE cDNA Amplification Kit (Clontech, USA) following the manufacturer’s protocol. The amplified products were TA-cloned into the pMD19-T vector (TaKaRa Bio Inc., Japan) and subjected to Sanger sequencing.

#### Northern Blotting

Total RNA was extracted from colon cancer and adjacent tissues or HCT116 cells using the TRIzol reagent (Invitrogen). Equal amounts of RNA (10-20 µg) were denatured by formaldehyde and subjected to agarose gel electrophoresis. The RNA was then transferred onto a nylon membrane and cross-linked by UV exposure for 1minute. The membrane was pre-hybridized in hybridization buffer at 68°C for 1 hour, followed by hybridization with the DIG-labeled probe specific to the target RNA at 68°C overnight. After washing, the membrane was incubated with anti-digoxigenin-AP (Roche) for signal detection.

#### Cell proliferation assays

Wild-type cells and clones transfected with siRNA oligos or lentiviruses expressing tsTE1 were seeded at the density of 2500 cells per 96-well. We measured the cell viability during the 5-day culture period using the Cell Counting Kit-8 (CCK8) (TargetMol) according to the manufacturer’s protocol. Additionally, we directly assessed the DNA synthesis using Cell-Light™ EdU Apollo-567 in vitro imaging kit. Briefly, HCT-116 cells were treated with 50 μmol/L EdU for 2 hours, fixed with 4% paraformaldehyde, and then stained with Apollo dye and Hoechst successively. Images were taken with the confocal microscope (LEICA, Germany) with the fluorescence quantified using Image J software.

#### TOP1 DNA supercoil relaxation

Referring to the procedure described by Baranello et al.^53^, we assessed the effect of tsTE1 on TOP1-mediated DNA supercoil relaxation using agarose gel electrophoresis. Briefly, recombinant human TOP1 protein (Sino Biological, China) was incubated with tsTE1 or LUC control RNA for 6 minuets in TOP1 buffer (10 mM Tris-HCl pH 7.5, 50 mM KCl, 5 mM MgCl2, 0.1 mM EDTA, 15 ug/ml BSA) on ice, followed by the incubation with 300ng plasmid DNA (pGL3) at 37°C for 20 minutes. The reaction was then terminated with TE-SDS 1% proteinase K (200 µg/ml) and the purified DNA products were purified and separated on a 1.4% (w/v) agarose gel using TAE buffer (pH 7.6). Supercoiled and relaxed plasmid DNA were semi-quantified using Image J software.

#### RNA pull down assay

Biotin-labeled tsTE1 RNAs were synthesized in vitro with T7 RNA polymerase (New England Biolabs, Inc.) and biotin-labeled UTP (Roche, Inc.). These tsTE1 RNAs were subsequently incubated with protein lysates of HCT-116 cell line for 4 hours at 4C allowing the formation of RNA-protein complexes. The RNA-protein mix was then incubated with streptavidin magnetic beads for 30 minutes at room temperature. The captured RNA-protein complexes were subsequently analyzed via SDS-PAGE followed by silver staining. Protein bands enriched in the tsTE1 lane compared to the control (LUC RNA) were excised and subjected to mass spectrometry to identify potentially interacting proteins. Immunoblotting was also employed to verify the interaction between tsTE1 and protein candidates. Truncated forms of TOP1 protein was designed based on the domains. The full-length and truncated DNA sequences of TOP1 were synthesized and then cloned into the pCMV-N-Flag vector.

#### Western blotting analysis

Proteins were subjected to SDS-PAGE and transferred to the nitrocellulose membrane. The membrane was blocked with 5% non-fat milk to prevent non-specific binding and then incubated with primary antibodies overnight at 4°C. Following washes, the membrane was next incubated with secondary antibodies for 1.5 hours at room temperature. Protein bands were detected using enhanced chemiluminescence (ECL) substrate and imaged with a chemiluminescence imaging system.

#### Xenograft in nude mice

To assess tthe impact of tsTE1 on tumor proliferation in vivo, we established a subcutaneous xenograft model using HCT-116 cells in nude mice. HCT-116 cells were digested, counted, and adjusted to 2×10C cells/200 μL in DMEM. Six-week-old male BALB/c-nu/nu mice (n=6/group) received subcutaneous injections of cell suspension. To overcome siRNA instability in vivo, LNA-modified antisense oligonucleotides (ASOs) were used for tsTE1 knockdown. ASO-tsTE1 was selected based on the efficiency and specificity in vitro. Mice received subcutaneous injections of ASO-tsTE1 (experimental) or ASO-Control. Tumor growth was monitored, with length (L) and width (W) measured every 3-4 days post-tumor formation. Tumor volume was calculated as (L×W²)/2. All procedures complied with guidelines approved by the Animal Care and Use Committee of the Fudan University Shanghai Cancer Center (FUSCC).

#### TEtrans RNA preparation for immunogenecity validation

The TAKARA-AGG-SP-MITD-EGFP-BSAI vector constructed in our recent work for mRNA therapeutic vaccines was employed as the in vitro transcription templetes^55^. The ORF sequences of TEtrans of interest were cloned into the plasmid with a 3×Flag tag added to the C-terminal. The recombinant plasmid was linearized by Hind III digestion and used as a template for in vitro transcription. The template was co-incubated with T7 RNA polymerase, nucleotides, 5′-cap analogues and in vitro transcription buffer for 2 hours in 37°C water bath. TEtrans RNAs were purified with RNA recovery column (ZYMO) after the DNase I treatment to remove the DNA template.

#### In vitro immunogenecity assay

PBMCs were isolated from healthy donors or patients with STAD using Human Lymphocyte Separation Tube (DaKeWe, China) according to the manufacturer’s instructions. Monocytes, B cells and CD8+ T cells were positively selected using CD14, CD19 and CD8 magnetic beads (STEMCELL Technologies) respectively. The culture procedures for immune cells were modified from the protocol established by Ali, M et al.^68^. Monocytes were cultured in AIM V medium (Gibco) supplemented with 1% inactivated human AB serum, 800 IU/mL recombinant human GM-CSF and 200 IU/mL IL-4 (Peprotech) for 5 days, inducing the differentiation into Mo-DCs which were subsequently maturated overnight with 100 IU/mL IFN-γ (Peprotech) and 10 ng/mL lipopolysaccharide (Sigma) for antigen presentation to naive CD8+ T cells. CD8+ T Cells were cultured in AIM V medium supplemented with 5% inactivated human AB serum, 5 ng/mL IL-7 and 5 ng/mL IL-15 (Peprotech) during the induction of antigen-specific clones. B cells were cultured in AIM V medium containing 8% inactivated human AB serum, 40 ng/mL IL-4 (Peprotech), 50 ng/mL IL-21 (Peprotech), 1 μg/mL CD40L (ACROBiosystems Group, China), and 50 μM β-mercaptoethanol for 7-10 days before used for mRNA transfection and antigen presentation.

For antigen loading, we performed mRNA transfection using the CALNP™ mRNA in vitro transfection reagent, which utilizes advanced RNA delivery technology for efficient and low-toxicity transfection. The TEtrans mRNA was diluted to 500 ng/μL, mixed with Reagent A and Reagent B in a volume ratio of 1:7:2 followed by the incubation of 10 minutes at room temperature. The mixture was then added to the cell culture medium at a 1:10 dilution for incubation over 24-48 hours. The transfection efficiency was assessed by the detection of Flag-tagged TEtrans-derived protein with western blotting.

To verify the existence of TEtrans antigen-sepcfic CD8+ T cell precursors in healthy donors, transfected Mo-DCs were co-incubated with CD8+ T cells at the ratio of 1:5 for 7-10 days. The culture medium was AIM V medium supplemented with 5% inactivated human AB serum, 5 ng/mL IL-7, 5 ng/mL IL-15 and 30ng/ml IL-21 (Peprotech). For the detection of antigen-specific CD8+ T cell clones in patients with gastric cancer, we levaraged the unfractionated PBMCs as antigen-presenting cells, transduced them with TEtrans RNA and then expanded T cells in an antigen-specific way referring to a published STAR protocol^69^. In either situations, the antigen-specific T cell responses were both detected with ELISPOT assay quantifying IFN-γ secretion after the restimulation with transfected or control B cells.

#### Tumor Cell Killing Assay

K562-HLA-A*02:01 cells were transfected with mRNA or not and stained with CFSE at different concentrations to distinguish transfected and control tumor cells. These labeled K562 cells mixed at the ratio of 1:1 (5×10^4^ each) and co-cultured with 2×10^5^ TEtrans-induced CD8+ T cells with the addition of 2 μg/mL CD28 antibody (BioGems) for overnight incubation. Subsequently, cell viability was assessed by flow cytometry, and the ratio of high to low CFSE-labeled viable cells was used to determine the specific killing ratio of T cells. The activation of CD8+ T cells was spontaneously evaluated by FACS detecting antigen-induced IFN-γ.

### Human research participants

PBMCs from two patients with gastric cancer were collected from Department of Medical Oncology of Fudan University Shanghai Cancer Center under a protocol approved by the Ethics Committee of Fudan University Shanghai Cancer Center. All patients in this study provided written informed consent for sample collection and data analysis.

### Statistical Analysis

All statistical analysis was performed in R software (v.4.1.0) or GraphPad Prism (v.8.0.2). Experimental data shown in column graphs represent meanC±Cstandard deviation. Detailed statistical tests can be found in the figure legends. Significance values: *PC<C0.05; **PC<C0.01; ***PC<C0.001; ns, no significant difference. TE class and subfamily enrichment analysis was performed with Fisher’s exact tests.

## Data Availability

The ASJA program is available at https://github.com/HuangLab-Fudan/ASJA. Publicly accessible datasets utilized in this study are described in detail within the “Materials and Methods” section. Original datasets and custom code generated during this work are available from the corresponding author upon reasonable request.

## Contributions

Conceptualization, Supervision and Writing - Review & Editing, S.H., X.Z. and X.H.; Data Curation, Software, Methodology, and Analysis, Z.L., Q.L., H.Z., P.L. and J.Z.; Visualization, Z.L. and Y.B.; Experimental Procedures, Z.L., Y.B., H.Y., Y.W. and Y.Y.; Resources, Z.H. and Y.L; Writing - Original Draft, Z.L. and Y.B.; All authors provided feedback on manuscript drafts.

## Supplementary figure legends

**Fig. S1: A.** TE splicing events identified in TCGA, CCLE, and GTEx cohorts. **B.** Comparison of the positional distribution of TE splicing events to all splicing events in TCGA. **C.** Over 90% of the splicing junctions within TEtrans are unannotated, a higher percentage compared to all TE-related splicing events. TE one_end and TE both_end refer to splicing sites where one end or both ends are provided by TEs, respectively. **D.** Distribution of the length and number of exons in TEtrans. **E.** TEtrans quantities of tumor and adjacent normal tissues in TCGA across various tumor types. **F.** Distribution of TEtrans quantities across different tissue types in the CCLE cell line collection. **G.** TEtrans are broadly expressed across various tumor types. The heatmap shows the expression frequencies of the top 30 most prevalent TEtrans in each tumor type. TEtrans prevalent in reproductive system cancers exhibit especially broad expression in pan-cancer level.

**Fig. S2: A.** Expression in tumor cell lines for two representative TEtrans transcripts shown in Figure 1D, validated by qPCR. **B.** Comparing the proportion of polyA tail signals located upstream of TTS for TEtrans, mRNA, and lncRNA, across 40bp and 150bp regions. “Total” represents the proportion of AAUAAA motifs detected, while “tp” refers to the true positive proportion after removing random background sequences containing polyA tail signals. The proportion within the 150bp range is similar between TEtrans and the other two mature transcript types. **C.** About 20.5% of TEtrans transcripts show the overlap with CAGE peaks curated by FANTOM project across TSS ± 500bp region. **D.** The IGV diagram illustrating the CAGE peaks near the TSS region of a typical TEtrans abundant in the MCF-7 cell line.

**Fig. S3: A.** Distribution of TE classes across four TEtrans splicing modes. **B.** Top: Density plots displaying donor and acceptor splice site distributions for the HERVK subfamily in One_Ele and One_subFam modes, showing a higher env splicing rate in One_subFam mode, similar to HERVH subfamily. Bottom: Density plots of donor and acceptor splice site distributions for the SVA_D element in One_Ele and Diff_Class modes. **C.** Schematic illustration for the splicing efficiency reporter vector. An 86 bp artificial intron, composed of conserved 5’ and 3’ splice site motifs, was inserted into the EGFP open reading frame of the pWPXL plasmid. The construct was transfected into 293T cells to assess the proportion of splicing products. **D.** Gel electrophoresis of qPCR products amplified from primers flanking the artificial intron in 293T cells. **E.** Quantification of splicing ratios following the insertion of various artificial introns.

**Fig. S4: A.** TEscore shows a significant positive correlation with tumor stemness features across multiple cancer types. **B.** Tumors with high TEscore exhibit higher chromosomal instability and intratumoral heterogeneity at a pan-cancer level. **C.** TEscore levels across molecular subtypes of upper gastrointestinal tumors. **D.** Correlation of TEscore with driver gene mutations in various cancer types. The heatmap color represents odds ratio wihch is the TEscore fold change between driver gene-mutated samples and wild-type counterparts. Fisher’s exact test, * for P < 0.05, ** for P < 0.01, *** for P < 0.001, blank boxes denote no significant difference. **E.** Association of TEscore with HLA class I allele loss of heterozygosity (HLA_LOH) events and tumor mutational burden (TMB).

**Fig. S5: A.** The tsTE1 expression in colorectal and liver cancer cell lines measured by qPCR. **B.** Agarose gel electrophoresis of 5’RACE and 3’RACE PCR products for tsTE1 transcript. Yellow arrows indicate the corresponding PCR products, with sequencing results shown to the right of the gel image. **C.** Northern Blot analysis of tsTE1 for total RNAs of HCT-116 and NCM460, confirming the size of full-length transcript amplified from RACE assays. **D.** Northern Blot analysis of tsTE1 for three pairs of colon cancer and adjacent normal tissues. **E.** High tsTE1 expression is significantly associated with worse progression-free survival in TCGA READ patients. P-value was calculated using the log-rank test.

**Fig. S6: A.** EdU assay to assess DNA synthesis in cells transfected with siRNA targeting tsTE1 compared to control. **B.** Quantification of EdU fluorescence intensity in tsTE1-knockdown and control groups. **C.** RNA pulldown assay using biotin-labeled sense and antisense strands of tsTE1. Silver-stained PAGE gel shows differential protein bands enriched in the tsTE1 sense strand pulldown (indicated by arrows). **D.** Predicted secondary structure of tsTE1 using RNAfold (http://rna.tbi.univie.ac.at/cgi-bin/RNAWebSuite/RNAfold.cgi) which guided the design of tsTE1 truncation mutants. **E.** Agarose gel electrophoresis of full-length and truncated tsTE1 constructs.

**Fig. S7: A.** Distribution of tumor-specific and tumor-associated splice sites of TEtrans coding candidates across cancer types. **B.** High TEtrans neoantigen load is associated with increased IFN-γ responsiveness, CD8+ T cell infiltration, activated dendritic cell (DC) populations, and immune checkpoint expression at the pan-cancer level. **C.** Length of transcripts and predicted open reading frames (ORFs) of TEtrans coding candidates. **D.** Exon numbers of TEtrans coding candidates. **E.** Mapping positions for TEtrans predicted ORFs. "Junction" indicates ORFs spanning splice sites, "Exon1" represents ORFs derived from the first exon, and "Other_exon" refers to ORFs derived from other exons.

**Fig. S8: A-C.** The tumor-specific expression of three additional candidate TEtrans: T2432 **(A)**, T4960 **(B)**, and T5739 **(C)** across various cancer types. **D.** Western blot analysis of 293T cells transfected with four candidate TEtrans, verifying the the predicted sizes of encoded proteins. **E.** Validation of mRNA transfection efficiency in human PBMCs, DCs, and CD19+ B cells using the CALNP™ RNA delivery system. Bright field (top) and green fluorescence images (bottom) are shown. **F-G.** IFN-γ ELISPOT spots quantificaton evaluating the induction of activated CD8+ T cells from two healthy donors by four candidate TEtrans. Due to limited cell numbers, each TEtrans group was repeated twice. **H.** IFN-γ ELISPOT spots quantification evaluating the T4960-specific activation of CD8+ T cell clones from two gastric cancer patients (P1 and P2). **I.** Schematic of the tumor cell killing assay. K562-HLA0201 cell lines transfected with TEtrans mRNA or not were labeled with different concentrations of CFSE, mixed in equal proportions, and co-cultured with antigen-specific CD8+ T cell clones. The ratio of live cells with different fluorescence intensities was measured by flow cytometry to calculate antigen-specific killing efficiency. **J.** T2432-induced CD8+ T cell clones specifically kill K562-HLA0201 cells expressing the corresponding antigen. **K.** Intracellular IFN-γ flow cytometry analysis showing the reactivation of T2432-induced CD8+ T cell clones by K562 cells expressing T2432.

**Fig. S9: A.** Experimental design for mRNA vaccination evaluating in vivo immunogenecity of mTEtrans. C57BL/6N mice were subcutaneously inoculated with 1×105 B16F10 cells and vaccinated with mRNA vaccines encoding mTEtrans_1(n=5) / combinations of 8 hotspot mutations(n=5) / control(n=5) on day 3 and 10. **B.** Representative flow cytometry images demonstrating the elevated proportion of mTEtrans_1-specfic CD8+ T cells in mouse spleenocyte after twice vaccinations.

